# Dynamics of Auxilin1 and GAK in clathrin-mediated traffic

**DOI:** 10.1101/718726

**Authors:** Kangmin He, Eli Song, Srigokul Upadhyayula, Song Dang, Raphael Gaudin, Wesley Skillern, Kevin Bu, Benjamin R. Capraro, Iris Rapoport, Ilja Kusters, Minghe Ma, Tom Kirchhausen

**Author notes:** These authors contributed equally to this work (Kangmin He, Eli Song, Srigokul Upadhyayula). Corresponding author: Dr. Tom Kirchhausen Harvard Medical School 200 Longwood Ave, Boston, MA 02115.

## Abstract

Clathrin coated vesicles formed at the plasma membrane lose their clathrin lattice within seconds of pinching off, through the action of the Hsc70 “uncoating ATPase”. The J-domain containing proteins, auxilin1 (Aux1) and auxilin2/cyclin-G dependent kinase (GAK), recruit Hsc70. Aux1 and GAK are closely related homologs, each with a phosphatase- and tensin-like (PTEN-like) domain, a clathrin-binding region, and a C-terminal J-domain; GAK has an additional, N-terminal Ser/Thr kinase domain. The PTEN-like domain has no phosphatase activity, but it can recognize phosphatidylinositol phosphate head groups. Aux1 and GAK appear on coated vesicles in successive transient bursts, immediately after dynamin mediated membrane scission has released the vesicle from the plasma membrane. We show here that these bursts represent recruitment of a very small number of auxilins such that even 4-6 molecules are sufficient to mediate uncoating. In contrast, we could not detect auxilins in abortive pits or at any time during coated-pit assembly. We have also shown previously that clathrin coated vesicles have a dynamic phosphoinositide landscape, and we have proposed that lipid head group recognition might determine the timing of Aux1 and GAK appearance. We now show that differential recruitment of Aux1 and GAK correlates with temporal variations in phosphoinositide composition, consistent with a lipid-switch timing mechanism.

## INTRODUCTION

Endocytic clathrin coats recruit molecular cargo as they assemble at the plasma membrane as coated pits and pinch off as coated vesicles. Cargo delivery then requires shedding of the clathrin lattice to liberate the enclosed vesicle (Kirchhausen et al., 2014). Disassembly of the coat, driven by the Hsc70 “uncoating ATPase” (Braell et al., 1984; Schlossman et al., 1984; Ungewickell, 1985), occurs just a few seconds after vesicle release (Lee et al., 2006; Massol et al., 2006); the timing of Hsc70 recruitment depends in turn on arrival of a J-domain containing protein, auxilin, immediately after the vesicle separates from the parent membrane (Lee et al., 2006; Massol et al., 2006). Human cells have two auxilin isoforms (Eisenberg and Greene, 2007). Auxilin2, expressed in all cells, has both a cyclin-G dependent kinase (GAK) domain and a phosphoinositide-phosphatase-like domain N-terminal to its clathrin-binding and J-domains. The latter domain, although catalytically inactive, is a phosphatase and tensin-like (PTEN) module (Guan et al., 2010). Auxilin1, expressed principally in neurons, has PTEN-like, clathrin-binding, and J-domains, but lacks the N-terminal kinase. We refer to these two auxilins as GAK and Aux1, respectively.

Aux1 and GAK contact three different clathrin heavy chains, from three different triskelions, when it associates with a clathrin coat (Fotin et al., 2004a). Because of the highly intertwined organization of a clathrin lattice (Fotin et al., 2004b), each of the contacts is with a distinct heavy-chain segment. The heavy chain also has an Hsc70 attachment site near its inward projecting C-terminus (Scheele et al., 2003), to which a nearby auxilin J domain can recruit Hsc70:ATP. A cryo-EM structure of Aux1- and Hsc70 bound coats at ∼11 Å resolution (Fotin et al., 2004a) suggested that no more than one Hsc70 could associate with an assembled trimer, since the three docking sites appeared too close to each other to allow multiple occupancy, but overlap of the three possible orientations precluded fitting the known Hsc70 structure to density. From *in vitro*, single-molecule analysis of Hsc70 driven uncoating we found that for coats saturated with auxilin, Hsc70 triggered rapid, apparently cooperative disassembly when it had accumulated to a critical level of about one molecule per two clathrin trimers (Bocking et al., 2011). Ensemble kinetic experiments from another laboratory (Rothnie et al., 2011) suggested a quite different picture, however, in which sequential binding (and concomitant ATP hydrolysis) by three Hsc70 molecules would be required to release each trimer.

In the work described here, we have sought to resolve these conflicting conclusions by studying the molecular mechanism of uncoating in the natural environment of a living cell. We have expressed, from its endogenous locus, Aux1 or GAK bearing a genetically encoded fluorescent tag and followed recruitment to endocytic coated vesicles by TIRF imaging with single-molecule sensitivity. We find that the burst-like recruitment of Aux1 or GAK that leads to uncoating, following scission of the membrane vesicle, is in all cases sub-stoichiometric and that uncoating with normal kinetics can occur after just 4-6 molecules of one or the other protein has accumulated. We further show that auxilins are absent from assembling pits, thus ruling out the possibility that earlier arrival could lead to Hsc70-driven clathrin exchange during coated-pit formation or to uncoating of an incomplete lattice and hence to a futile assembly-disassembly cycle.

Continuous lipid modification provides a potential mechanism by which auxilin could detect that a vesicle has separated from the parent membrane (He et al., 2017). Proposals for the mechanism by which the uncoating machinery distinguishes a pinched-off vesicle from maturing coated pit have invoked phosphoinositide recognition by PTEN-like domain and an enzymatic mechanism that alters vesicle lipid composition following budding from the parent membrane (Cremona et al., 1999; He et al., 2017). We showed recently that the phosphoinositide composition of an endocytic coated vesicle remains unchanged until the moment of separation from the plasma membrane but then undergoes a well-defined series of sequential modifications, and we identified enzymes responsible for some of the transformations. We have now determined, with single-molecule sensitivity, the correlation of arrival times and quantities of Aux1 and GAK with steps in coat formation and disassembly. Recruitment of Aux1 and GAK then follows the temporal variations in phosphoinositide composition, dictated by the differential specificities of their PTEN-like domains. We further show that recruitment of only a small number of auxilin molecules is enough for complete uncoating. These observations define a coincidence-detection and lipid-switch timing mechanism that distinguishes a coated vesicle from a coated pit and that launches the uncoating process as soon as coated-vesicle formation is complete.

## RESULTS

### Dynamics of auxilin mediated uncoating in gene-edited cells expressing fluorescently tagged auxilins

To study uncoating-associated recruitment of Aux1 and GAK, we established cell lines expressing fluorescently tagged Aux1 or GAK by homozygous replacement with a corresponding chimera bearing EGFP at its N-terminus (EGFP-Aux1 or EGFP-GAK) (Fig. 1a and Supplementary Fig. 1a-c). The same cells also had either full replacement of clathrin light chain A (CLTA) with the fluorescent chimera CLTA-TagRFP or full replacement of AP2-σ2 with AP2-σ2-TagRFP. We chose SUM159 cells (Forozan et al., 1999), a largely diploid, human breast cancer–derived cell line because, like HeLa and other non-neuronal lines (Borner et al., 2012; Hirst et al., 2008), these cells express both Aux1 and GAK (Supplementary Fig. 1b,c). We confirmed that the clathrin-mediated endocytic efficiency of the gene-edited cells was similar to that of the parental cells, by using flow cytometry to measure receptor-mediated uptake of fluorescently tagged transferrin (Supplementary Fig. 1d,e).

**Figure 1.**
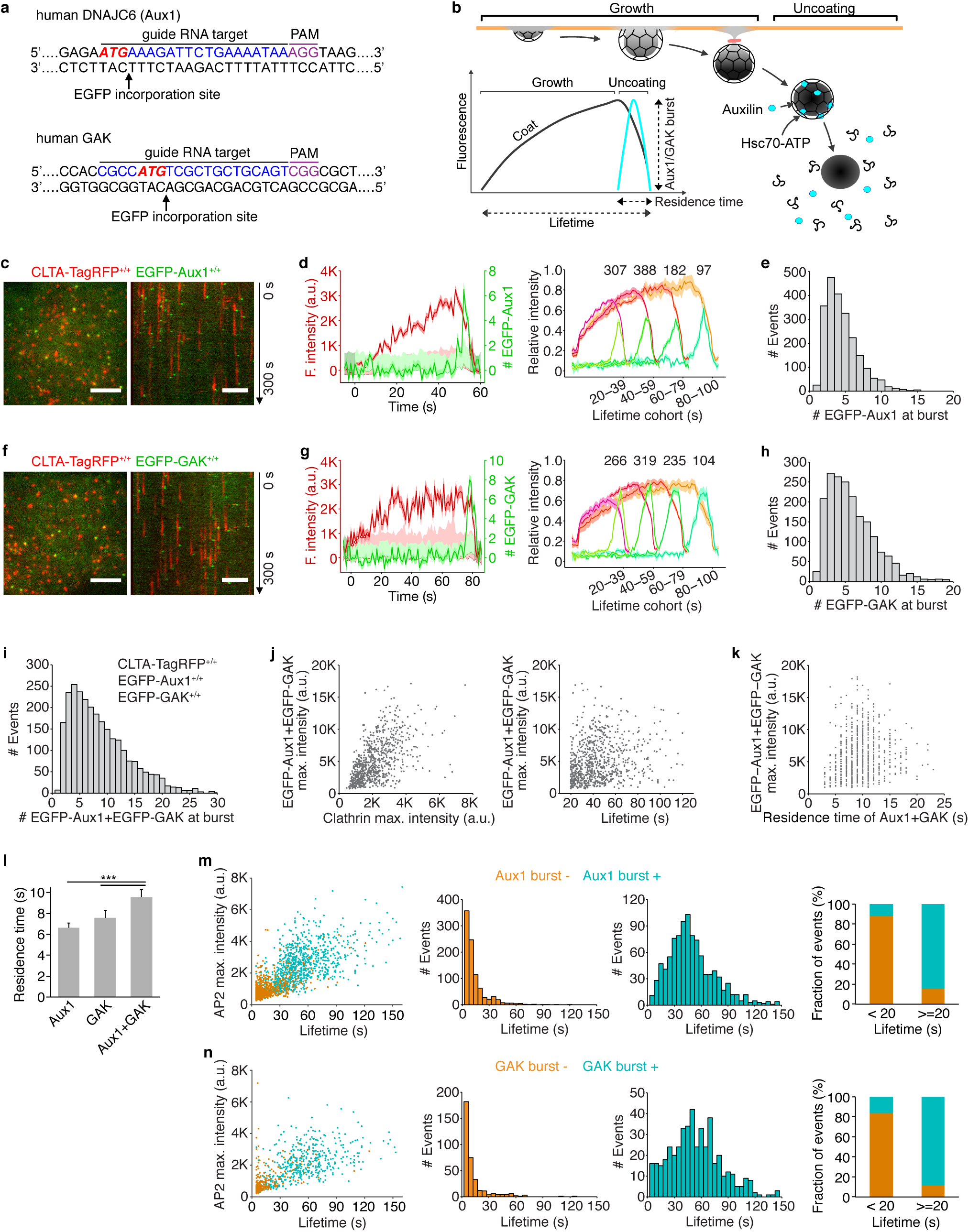
Recruitment of Aux1 and GAK to clathrin-coated vesicles in genome-edited cells. (**a**) CRISPR/Cas9 gene-editing strategy used to incorporate EGFP at the N-terminus of Aux1 or GAK. The target sequence at the genomic locus of gene *DNAJC6* (Aux1) recognized by the single-guide RNA is highlighted. The protospacer adjacent motif (PAM), the start codon ATG (red) and the site of EGFP incorporation upon homologous recombination are highlighted. (**b**) Schematic representation of the bursts of Aux1/GAK during uncoating of clathrin-coated vesicles (modified from Massol et al., 2006). (**c**) Snapshot (left) and kymograph (right) from a representative TIRF microscopy time series showing transient burst recruitment of EGFP-Aux1 in coated vesicles containing CLTA-TagRFP in SUM159 cell double gene-edited for EGFP-Aux1^+/+^ and CLTA-TagRFP^+/+^. Time series with single molecule detection sensitivity for EGFP acquired for 300 s at 1-s intervals using 100 ms exposures. The kymograph was shifted laterally by 5 pixels. Scale bars, 5 μm. (**d**) Left panel: Representative plot of a single endocytic event showing fluorescence intensity traces for CLTA-TagRFP and EGFP-Aux1 (arbitrary units for CLTA; number of molecules for Aux1) together with estimated uncertainties (one S.D., dark shade), corresponding local backgrounds (thin lines), and significance threshold above background (∼2 S.D., light shade). Right panel: Averaged fluorescence intensity traces (mean ± S.E.) for CLTA-TagRFP and EGFP-Aux1 from endocytic clathrin-coated pits and vesicles automatically identified in eight cells and then grouped in cohorts according to lifetimes. The numbers of traces analyzed are shown above each cohort. (**e**) Distribution of the maximum number of EGFP-Aux1 molecules recruited during the uncoating burst (2198 traces from 17 cells). (**f-h**) Transient burst of EGFP-GAK in coated vesicles containing CLTA-TagRFP in SUM159 cell double gene-edited for EGFP-GAK^+/+^ and CLTA-TagRFP^+/+^. Cohorts in **g** are from eight cells; distributions in **h** from 16 cells. (**i**) Distribution of the maximum number of EGFP-Aux1 together with EGFP-GAK molecules recruited during uncoating of clathrin-coated vesicles (2636 traces) from 31 cells triple-edited for EGFP-Aux1^+/+^, EGFP-GAK^+/+^ and CLTA-TagRFP^+/+^ (**j**) Scatter plots for the maximum fluorescence intensities of EGFP-Aux1 and EGFP-GAK (781 events) as a function of the maximum fluorescence intensity of CLTA-TagRFP (left) (Pearson correlation coefficient *r* = 0.569) or lifetime of clathrin-TagRFP (right) (Pearson correlation coefficient *r* = 0.212) from nine cells triple-edited for EGFP-Aux1^+/+^, EGFP-GAK^+/+^ and CLTA-TagRFP^+/+^. (**k**) Scatter plots for the maximum fluorescence intensities of EGFP-Aux1 and EGFP-GAK (716 events) as a function of the duration of Aux1 and GAK bursts (Pearson correlation coefficient *r* = 0.132) from nine cells triple-edited for EGFP-Aux1^+/+^, EGFP-GAK^+/+^ and CLTA-TagRFP^+/+^. (**l**) Duration of Aux1 and GAK bursts during uncoating of coated vesicles in cells gene-edited for EGFP-Aux1^+/+^ (six cells, Aux1), EGFP-GAK^+/+^ (five cells, GAK), or EGFP-Aux1^+/+^ and EGFP-GAK^+/+^ (nine cells, Aux1+GAK) together with CLTA-TagRFP^+/+^; \*\*\**P* < 0.001 by one-way ANOVA with Tukey’s comparison test. (**m**) Left panel: Scatter plot for the maximum fluorescence intensities and lifetimes from AP2-TagRFP tracks with (996 traces) or without (919 traces) detectable EGFP-Aux1 burst, from nine cells double gene-edited for EGFP-Aux1^+/+^ and AP2-TagRFP^+/+^. Middle panels: lifetime distributions from AP2-TagRFP tracks without or with detectable EGFP-Aux1 burst. Right panel: fraction of AP2 tracks of lifetime shorter or longer than 20 s with or without detectable EGFP-Aux1 burst. (**n**) Left panel: Scatter plot for the maximum fluorescence intensities and lifetimes of AP2-TagRFP tracks with (467 traces) or without (350 traces) detectable EGFP-GAK burst, from six cells double gene-edited for EGFP-GAK^+/+^ and AP2-TagRFP^+/+^. Middle panels: lifetime distributions from AP2-TagRFP tracks without or with detectable EGFP-GAK burst. Right panel: fraction of AP2 tracks of lifetime shorter or longer than 20 s with or without a detectable EGFP-GAK burst.

We confirmed the burst-like recruitment of EGFP-Aux1 and EGFP-GAK, restricted to the time of clathrin uncoating (Fig. 1b-h), by analyzing fluorescent traces from time series acquired by TIRF microscopy of gene-edited cells (see Methods). The Aux1 bursts and most GAK bursts occurred at the relatively immobile clathrin spots we have shown to be associated with endocytic events (Ehrlich et al., 2004). GAK, but not Aux1, also associates with the more mobile, clathrin-coated structures emanating from the trans-Golgi network (TGN) and endosomes (Greener et al., 2000; Kametaka et al., 2007; Lee et al., 2005; Zhang et al., 2005), and a few objects in the EGFP-GAK expressing cells indeed appeared mobile in our TIRF microscopy time series. We confirmed this differential recruitment by full volume 3D live-cell lattice light-sheet microscopy (Fig. 2a,b,d).

**Figure 2.**
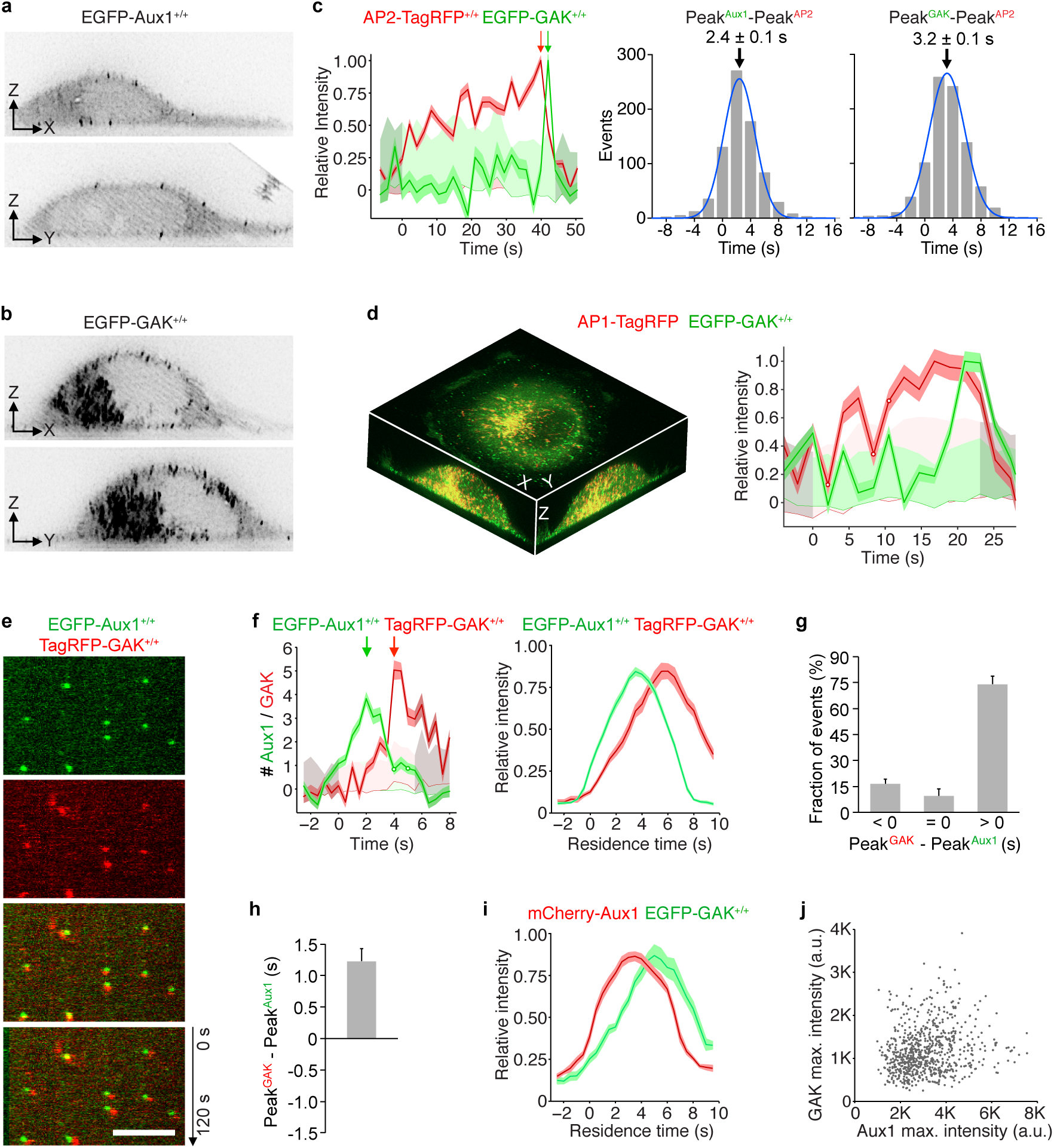
Temporal and spatial distributions of Aux1 and GAK. (**a**) Maximum intensity projections from a thin 2 μm optical section of a gene-edited cell expressing EGFP-Aux1 recorded in 3D by lattice light-sheet microscopy (**b**) Maximum intensity projections from a thin 2 μm optical section of a gene-edited cell expressing EGFP-GAK recorded in 3D by lattice light-sheet microscopy. (**c**) Representative plot from 3D automated trackings of AP2-TagRFP and EGFP-Aux1 (872 traces from six cells), or AP2-TagRFP and EGFP-GAK (755 traces from six cells) in the double gene-edited cells imaged by lattice light-sheet microscopy. Distribution (fit with a single Gaussian) of the interval between maximum fluorescent signals for AP2 and Aux1 or AP2 and GAK (2.4 ± 0.1 s and 3.2 ± 0.1 s, mean ± S.E., respectively). (**d**) Distribution of GAK in cells gene-edited for EGFP-GAK^+/+^ and stably expressing AP1-TagRFP recorded in 3D by lattice light-sheet microscopy. Left panel: Maximum intensity projections in X-Y, X-Z and Y-Z of EGFP-GAK and AP1-TagRFP from a 3D rendered cell. Right panel: Representative plot of EGFP-GAK recruitment to a single AP1-positive carrier. (**e**) Snapshot from a representative TIRF microscopy time series showing the transient burst recruitment of EGFP-Aux1 and TagRFP-GAK^+/+^ on the plasma membrane of a cell double gene-edited for EGFP-Aux1^+/+^ and TagRFP-GAK^+/+^. Time series with single molecule detection sensitivity for EGFP and TagRFP acquired for 120 s at 1 s intervals using 100 ms exposures. Kymograph (bottom panel) shifted laterally by 5 pixels. Scale bars, 5 μm. (**f**) Representative plot (left panel) for a single endocytic event showing sequential recruitment of EGFP-Aux1 and TagRFP-GAK (recruitment peaks highlighted by arrows) imaged at 0.5 s intervals by TIRF microscopy. The traces (right panel) are averaged relative fluorescence intensity (mean ± S.E.) of EGFP-Aux1 and TagRFP-GAK for the cohort of EGFP-Aux1 burst with residence times lasting between three and 12 s (1516 traces from eight cells). (**g**) The relative timing differences (< 0 s, = 0 s and > 0 s) between peaks of EGFP-Aux1 and TagRFP-GAK recruitment (mean ± S.D., n = eight cells). (**h**) Distribution of interval between peaks of EGFP-Aux1 and TagRFP-GAK recruitment. The mean interval and S.D. from eight cells was 1.34 ± 0.26 s. (**i**) Averaged relative fluorescence intensity (mean ± S.E.) for bursts of duration between three and 12 s of transiently expressed mCherry-Aux1 and gene edited EGFP-GAK (1859 traces from eight cells). (**j**) Scatter plot for the maximum fluorescence intensities of EGFP-Aux1 and TagRFP-GAK (750 traces from eight cells double gene-edited for EGFP-Aux1^+/+^ and TagRFP-GAK^+/+^, Pearson correlation coefficient *r* = 0.189).

In cells expressing AP2-σ2-TagRFP, nearly all (∼90%) AP2-containing structures with lifetimes shorter than 20 s incorporated relatively small amounts of AP2, failed to recruit EGFP-Aux1 or EGFP-GAK and had a distinct quasi-exponential decay distribution of lifetimes associated with a stochastic coat dissociation process (Fig. 1m,n). These correspond to early abortive coated pits described in earlier work (Aguet et al., 2013; Ehrlich et al., 2004; Loerke et al., 2009). These characteristics match the properties of abortive coated pits described by Aguet et al using the dynamin burst as a surrogate marker (Aguet et al., 2013).

As we and others have shown (Aguet et al., 2013; Ehrlich et al., 2004; Hong et al., 2015; Loerke et al., 2009), the interval between initiation of an AP2-containing endocytic coated pit and it’s pinching off from the plasma membrane as a coated vesicle ranges between 20 and 150 s. Most of these structures (∼90%) incorporated greater amounts of AP2 than did the short-lived ones, displayed a multimode lifetime distribution characteristic of a process governed by the superposition of multiple steps and showed at the time of uncoating a burst of EGFP-Aux1 or EGFP-GAK (Fig. 1m,n). The multimode lifetime distribution is a signature of productive coated pits, precisely as defined by (Aguet et al., 2013; Ehrlich et al., 2004; Loerke et al., 2009). Those few longer-lived structures (∼10%) that failed to recruit auxilins had a characteristic quasi-exponential decay in their lifetime distributions (Fig. 1m,n) and probably corresponded to the late abortives observed previously (Aguet et al., 2013; Ehrlich et al., 2004; Loerke et al., 2009). These characteristics also match the properties of abortive coated pits using dynamin as a surrogate marker (Aguet et al., 2013; Ehrlich et al., 2004; Loerke et al., 2009). We inferred from these observations that most endocytic clathrin coated vesicles recruited both auxilins, and we confirmed this inference (as described below) by observing concurrent recruitment of EGFP-Aux1 and TagRFP-GAK in double edited SUM159 cells (Fig. 2e).

### Auxilins are not recruited to assembling clathrin-coated pits

We were unable to detect EGFP-Aux1 or EGFP-GAK recruitment while coated pits were assembling, even with the single molecule sensitivity of our TIRF microscopy (Fig. 1c,d,f,g; Supplementary Fig. 2 and Supplementary Video 1 and 2), and observed the burst recruitment only when assembly was complete. These results imply that an Aux1- and GAK-mediated process (and by inference Hsc70 activity) cannot account for published *in vivo* observations of partial exchange of clathrin between assembling endocytic coated pits and a cytosolic clathrin pool (Eisenberg and Greene, 2007; Wu et al., 2001). We note that the lattice of the assembling coat is competent to bind Aux1, since Aux1-based sensors for phosphatidylinositol-4-5-phosphate (PtdIns(4,5)P_2_), the predominant lipid species in the plasma membrane, appear at coated pits in quantities that follow the clathrin content (He et al., 2017). The observed exchange is presumably a consequence of the dynamic equilibrium present at the edge of any growing two-dimensional array. This mechanism is also consistent with our observation that abortive-pit disassembly did not require the auxilin-dependent uncoating machinery. We conclude that until a coat is complete, clathrin can dissociate from an exposed edge unless stabilized by interaction with other components.

### Recruitment specificity of auxilins

To investigate the mechanism responsible for the recruitment specificity, we first analyzed the burst dynamics of Aux1 and GAK by 3D tracking of EGFP-Aux1 or EGFP-GAK from cells gene-edited to express AP2-σ2-TagRFP together with EGFP-Aux1 or EGFP-GAK (Fig. 2c). The results, from time series obtained by 3D live-cell lattice light-sheet microscopy, showed that the time points for peak recruitment of Aux1 preceded those for GAK by ∼1 s (∼2.4 s and ∼3.2 s peak recruitment after initiation of uncoating, respectively) (Fig. 2c). We found the same differential timing in gene-edited cells expressing both Aux1 and GAK, labeled with different fluorescent tags, EGFP and TagRFP, respectively (Fig. 2e-h; Supplementary Fig. 3a,c; Supplementary Video 3). We minimized the likelihood that the observed recruitment delays were due to the fluorescent tags by showing that Aux1 and GAK maintained their differential timing in SUM159 cells gene-edited to express EGFP-GAK together with transient expression of mCherry-Aux1 or mCherry-GAK (Fig. 2i and Supplementary Fig. 3b,d). Moreover, we found the same relative recruitment dynamics of Aux1 and GAK in monkey COS-7 cells and human HeLa cells transiently expressing EGFP-Aux1 and mCherry-GAK (Supplementary Fig. 3e,f). Inspection of the time series illustrated in Fig. 2 showed no strong correlation between the maximum recruitment amplitudes of Aux1 and GAK into the same coated vesicle (Fig. 2j).

Why does Aux1 arrival precede recruitment of GAK? Our recent study (He et al., 2017) of phosphoinositide dynamics in endocytic compartments described sequential bursts of Aux1-based PtdIns(3)P and PtdIns(4)P sensors, ∼1−2 s apart, accompanying endocytic clathrin coated vesicle uncoating (He et al., 2017). The results in Fig. 3 show a close correspondence between the arrival times at endocytic coated vesicles of a Aux1-based PtdIns(3)P sensor and Aux1 and between the (∼1-2 s later) arrival times of a Aux1-based PtdIns(4)P sensor and GAK. Likewise, replacement of Aux1 with the unrelated Epsin1 (binds clathrin) used to generate another set of PtdIns(3)P and PtdIns(4)P sensors (He et al., 2017) also led to sequential burst arrivals ∼1-2 s apart (Supplementary Fig. 3h). These correlations suggest that phosphoinositide conversion determines the differential recruitment of Aux1 and GAK. This inference is also consistent with the results of *in vitro* lipid-protein overlay assays, which showed that PtdIns(3)P interacts preferentially with Aux1 (Massol et al., 2006) while PtdIns(4)P favors the PTEN-like domain of GAK (Lee et al., 2006). The observed lipid dependence of the Aux1- or GAK-mediated uncoating reaction was also observed in the *in vitro* single-object uncoating experiments described below using synthetic coated vesicles as substrate for the uncoating reaction.

**Figure 3.**
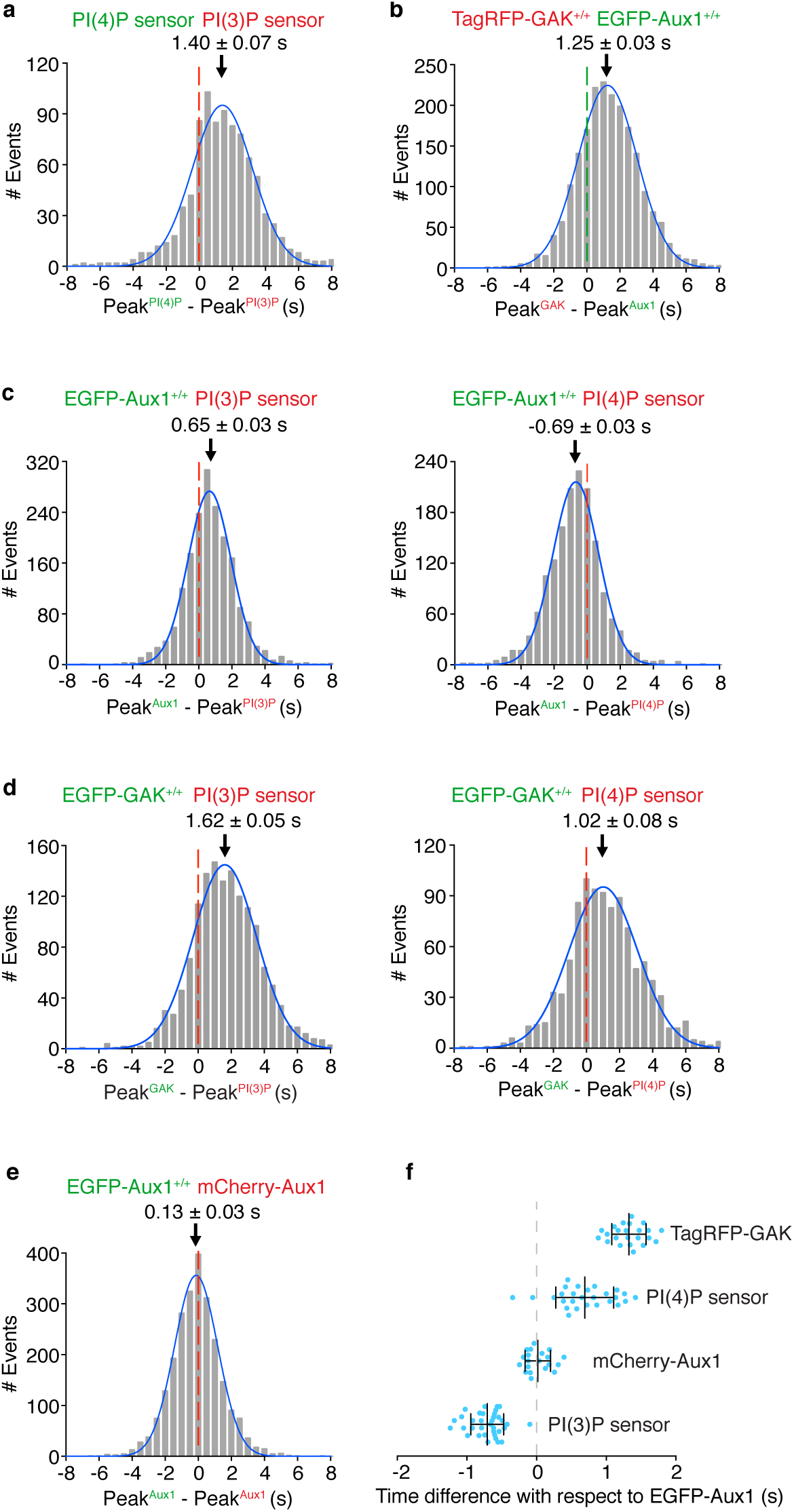
Comparison of recruitment times of Aux1, GAK and phosphoinositide sensors to endocytic clathrin-coated vesicles. Bottom (adherent) surfaces of cells transiently expressing various combinations of Aux1, GAK and Aux1-based PtdIns(3)P and PtdIns(4)P sensors imaged by TIRF microscopy every 0.5 s for 100 s. (**a**) Transient co-expression of the Aux1-based PtdIns(4)P (EGFP-P4M(DrrA)-Aux1) and PtdIns(3)P (mCherry-2xFYVE(Hrs)-Aux1) sensors in parental SUM159 cells. Distribution (fit with a single Gaussian) for the interval between the peaks within single events showing that the PtdIns(3)P sensor precedes the PtdIns(4)P sensor by 1.40 ± 0.07 s (mean ± S.E., 916 traces from 34 cells). (**b**) Distribution for the interval between the peaks within single events of EGFP-Aux1 and TagRFP-GAK in cells double gene-edited for EGFP-Aux1^+/+^ and TagRFP-GAK^+/+^ (−1.25 ± 0.03 s, mean ± S.E., 2033 traces from 23 cells). (**c**) Transient expression of PtdIns(3)P (mCherry-2xFYVE (Hrs)-Aux1) or PtdIns(4)P (mCherry-P4M (DrrA)-Aux1) sensor in gene-edited EGFP-Aux1^+/+^ cells. Distributions for the interval between burst peaks for Aux1 and the phosphoinositide sensors in the same event. Aux1 and PtdIns(3)P sensor: 0.65 ± 0.03 s, mean ± S.E., 1863 traces in 35 cells; Aux1 and PtdIns(4)P sensor: −0.69 ± 0.03 s; 1570 traces in 27 cells. (**d**) Transient expression of PtdIns(3)P (mCherry-2xFYVE (Hrs)-Aux1) or PtdIns(4)P (mCherry-P4M (DrrA)-Aux1) sensors in gene-edited EGFP-GAK^+/+^ cells. Distributions for the interval between burst peaks for GAK and the phosphoinositide sensors in the same event. GAK and PtdIns(3)P sensor (1.62 ± 0.05 s; 1435 traces in 36 cells); GAK and PtdIns(4)P sensor (1.02 ± 0.08 s; 1020 traces in 34 cells). (**e**) Transient expression of mCherry-Aux1 in gene-edited EGFP-Aux1^+/+^ cells. Distributions for the interval between burst peaks for mCherry-Aux1 and EGFP-Aux1 in the same event (0.13 ± 0.03 s; 2435 traces in 34 cells). (**f**) Interval between burst peaks of Aux1 in gene edited EGFP-Aux1^+/+^ cells and of gene edited TagRFP-GAK^+/+^ (n = 23 cells), or of transiently expressed PtdIns(4)P sensor (mCherry-P4M(DrrA)-Aux1, n = 27 cells), of mCherry-Aux1 (n = 34 cells) or of PtdIns(3)P sensor (mCherry-2xFYVE(Hrs)-Aux1, n = 35 cells), respectively, where the value of each spot represents the average (mean ± S.D) of the measurement obtained for a given single cell. The timing differences between the bursts for each group were statistically significant (*P*<0.001 by one-way ANOVA with Tukey’s comparison test).

To determine the importance of the PTEN-like domain for the specificity of intracellular targeting of auxilin, we swapped the PTEN-like domains of Aux1 and GAK and followed the intracellular location of transiently expressed chimeric variants (Fig. 4). As confirmed above (Fig. 2b,d), wild-type GAK appears in the perinuclear TGN and recycling endosomes, both of which are enriched in PtdIns(4)P (Kural et al., 2012; Wang et al., 2003), as well as in endocytic coated vesicles (Fig. 4a). As shown previously (Guan et al., 2010; Lee et al., 2006; Massol et al., 2006), the PTEN-like domain was essential for efficient targeting Aux1 or GAK to endocytic coated vesicles (Fig. 4c,h). A GAK chimera containing the PTEN-like domain of Aux1 replacing its own appeared exclusively in endocytic coated vesicles at the plasma membrane (Fig. 4e,f), and the extent to which this GAK-Aux1 chimera was recruited to those coated vesicles was similar to the extent of recruitment of wild-type Aux1 (Fig. 4g) but slightly less than that of wild-type GAK (Fig. 4a); the arrival time of this chimera also corresponded to the arrival time for wild-type Aux1 (Fig. 4m). The converse chimera, Aux1 with the PTEN-like domain of GAK, acquired the plasma membrane recruitment dynamics of wild-type GAK (Fig. 4j,k). Although the kinase domain was not required for recruiting GAK to endocytic coated vesicles, its presence substantially enhanced perinuclear targeting (Fig 4a,b). The kinase also enhanced perinuclear targeting of chimeric Aux1 with the GAK PTEN-like domain (Fig. 4j,k); adding it to wild-type Aux1 had no effect (Fig. 4l).

**Figure 4.**
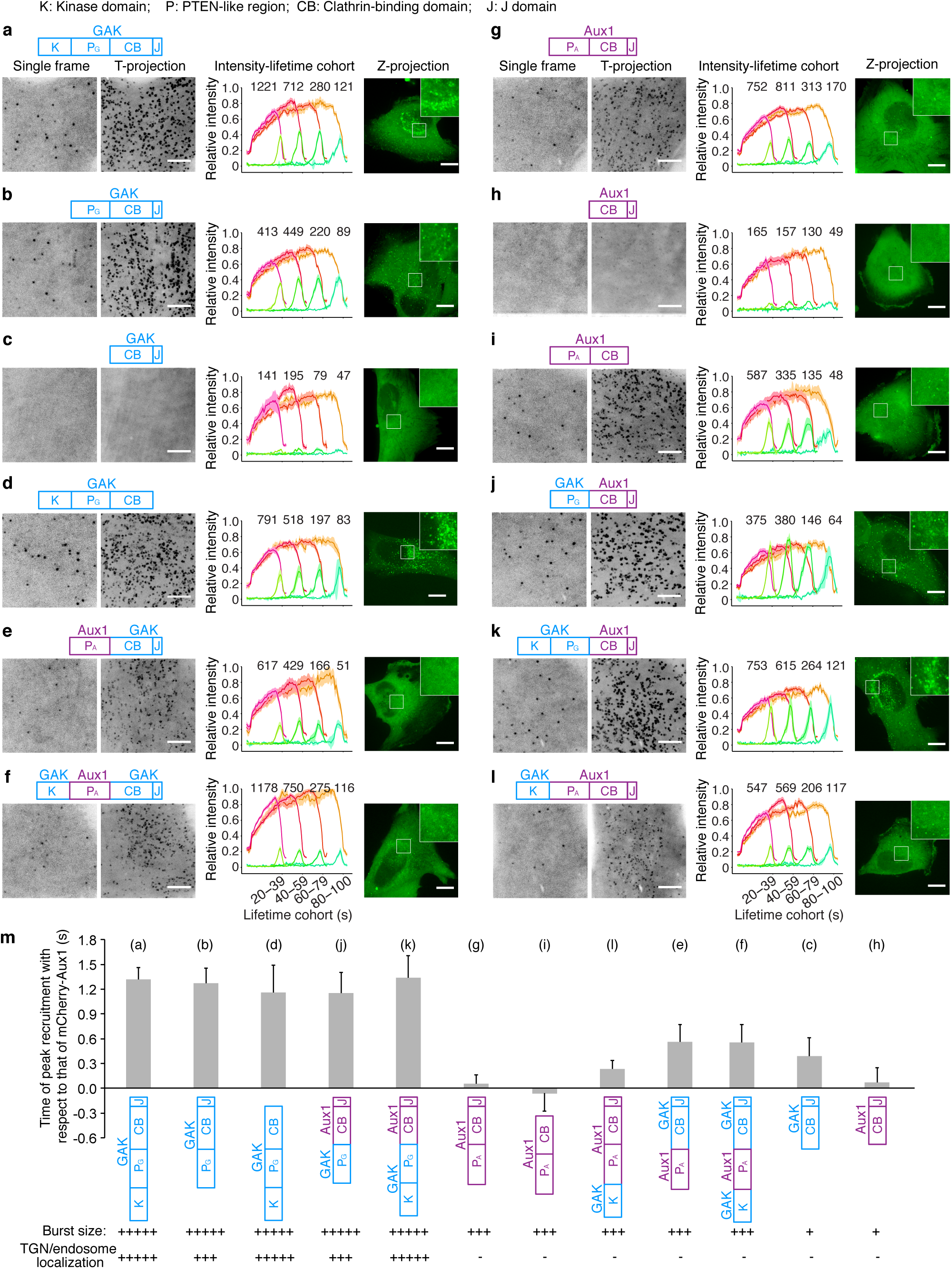
PTEN-like domains influence the recruitment of Aux1 and GAK. Bottom surfaces of gene-edited CLTA-TagRFP^+/+^ cells transiently expressing indicated EGFP-tagged constructs of Aux1 or GAK, imaged by TIRF microscopy every one s for 300 s; traces analyzed by automated 2D tracking. (**a-l**) Each panel shows a schematic representation indicating the domain organization of the construct: K, kinase domain of GAK (amino acid residues 1-347); P_A_ or P_G_, PTEN-like domain of GAK (360-766) or of Aux1 (1-419); CB, clathrin-binding domain of GAK (767-1222) or of Aux1 (420-814); J, J domain of GAK (1223-1311) or of Aux1 (815-910); a representative single frame and the corresponding maximum intensity time projection of the time series as a function of time (T-projection). The plots show averaged fluorescence intensity traces (mean ± S.E.) of CLTA-TagRFP (red) and EGFP-fused constructs (green), from 6-13 cells per condition, grouped by cohorts according to lifetimes. The numbers of analyzed traces are shown above each cohort. The cells were also imaged in 3D by spinning-disk confocal microscopy; the images at the right of each panel show representative maximum intensity z projections (Z-projection) from 34 sequential optical sections spaced 0.3 μm and include an enlarged region. Scale bars, 10 μm. (**m**) Mean interval between the fluorescence maxima for the indicated EGFP-fused construct (diagram immediately below plot) and the mCherry-Aux1 burst (mean ± S.D., n = 8-14 cells), in cells imaged at 0.5 s intervals for 60 s by TIRF microscopy. Below the construct schematics are qualitative estimates of the relative maximum amplitudes of fluorescence for the bursts at the plasma membrane and in regions of the TGN/endosome.

### Very few auxilins are sufficient to trigger uncoating

Previous *in vitro* ensemble studies have suggested that less than one auxilin per vertex is sufficient to elicit Hsc70-driven disassembly of synthetic coats (Bocking et al., 2011). To determine the requirements for Aux1 and GAK *in vivo*, we took advantage of the high sensitivity of TIRF microscopy to follow recruitment of EGFP-Aux1 or EGFP-GAK to endocytic coated vesicles in gene-edited cells. We used oblique illumination to record 3-5 min time series of frames recorded once per second. We calibrated the fluorescence intensity as described in our previous work (He et al., 2017). The duration of the EGFP-Aux1 or EGFP-GAK bursts (6-8 sec) was the same as previously observed for the corresponding ectopically expressed proteins (He et al., 2017; Lee et al., 2006; Massol et al., 2006) (Fig. 1l). Most bursts contained between 2-8 and 2-12 molecules with peak values of 3 +/- 1 and 4 +/- 2 molecules, for Aux1 and GAK respectively (Fig. 1e,h). Detailed analysis of 567 rapidly acquired EGFP-Aux1 or 276 EGFP-GAK traces revealed that the first recorded events were consecutive stepwise increases in fluorescence intensity (Supplementary Fig. 2e,f, selected examples). The number of Aux1 or GAK molecules recruited during each of the first two consecutive steps presented as histogram plots was determined by fitting the intensity distributions with the single-molecule calibration of fluorescence intensity (Supplementary Fig. 2e,f). The analysis suggests that recruitment begins with preferential arrival of a single auxilin during the initial step of recruitment followed by a second one (within the 62.5 ms time resolution of our measurements). The peak number of total auxilin molecules recruited during the burst ranged between 2-20 as determined by the peak fluorescence intensity of EGFP-Aux1 and EGFP-GAK simultaneously replaced in the same gene-edited cells (Fig. 1i). The duration of their combined bursts was slightly longer (∼10 sec) than each one alone (Fig. 1k,l). We found no correlation between the number of recruited auxilins and the observed uncoating rate, nor did the peak level correlate with the size of the coat, as estimated from the peak clathrin light-chain fluorescence intensity (Fig. 1j and Supplementary Fig. 1f,g). Most endocytic coated vesicle have between 36 and ∼100 vertices; our results show that uncoating proceeds even when only a relatively small proportion of the vertices have acquired an auxilin. We found a similarly sub-stoichiometric GAK occupancy of AP1-containing carriers; the amplitude of the GAK burst at perinuclear clathrin spots ranged from ∼5 to ∼25 (Supplementary Fig. 3g).

The experiments just described showed that when the two auxilins were present together, a relatively small number of Aux1 and GAK molecules were sufficient for normal uncoating. We then proceeded to establish the contribution of Aux1 or GAK alone to clathrin-mediated uptake of transferrin and to the kinetics of clathrin uncoating, by imaging cells lacking GAK (by CRISPR/Cas9-mediated knockout) or depleted of Aux1 (by shRNA). Complete elimination of GAK (Supplementary Fig. 4a) had no significant effect on transferrin uptake (Supplementary Fig. 4b), while slightly increasing the lifetimes of clathrin coated structures at the plasma membrane and the number of EGFP-Aux1 molecules recruited during the burst (Fig. 5a). The expression level of Aux1 was unaffected (Supplementary Fig. 4a). The interval associated with the EGFP-Aux1 burst (proportional to the amount of time required to uncoat) also increased slightly (Supplementary Fig. 4c). We obtained similar results with a transient, shRNA-based depletion of GAK (Supplementary Fig. 4d-g). Because we were unable to eliminate Aux1 by CRISPR/Cas9-mediated knockout, we used transient depletion with shRNA (Supplementary Fig. 4h). Aux1 depletion barely affected the rate of transferrin uptake (Supplementary Fig. 4i) or the efficiency of uncoating (Fig. 5b and Supplementary Fig. 4j) but led to a small increase in the peak number of GAK molecules recruited during the burst at the time of uncoating (Fig. 5b), mirroring the effect of GAK elimination.

**Figure 5.**
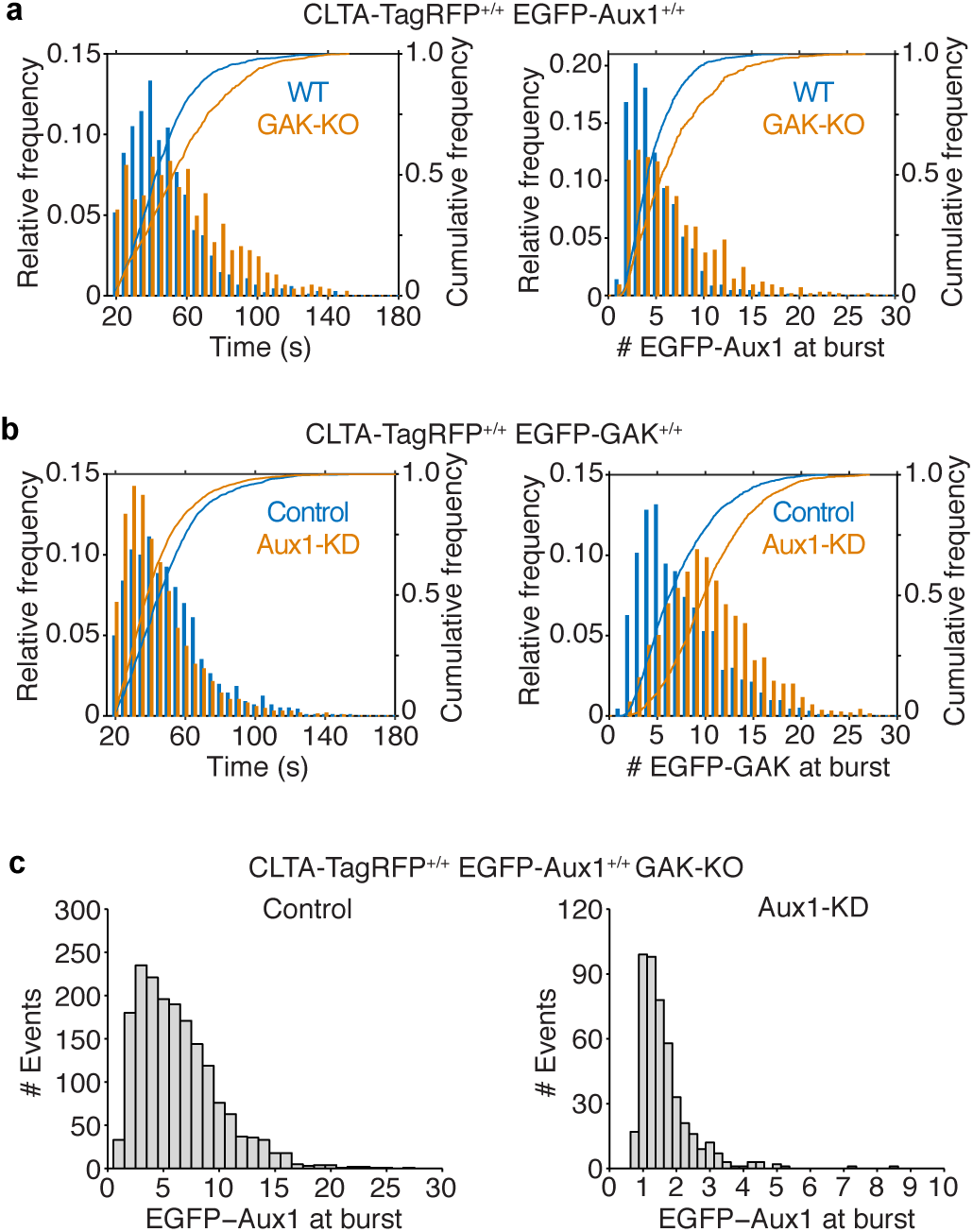
Numbers of auxilin molecules during uncoating. (**a**) Effect of GAK knockout on recruitment of Aux1 to endocytic clathrin coated vesicles. GAK was eliminated by CRISPR/Cas9-targeted knockout in cells gene-edited for EGFP-Aux1^+/+^ and CLTA-TagRFP^+/+^. Histogram and cumulative distributions showing significant increases in the number of EGFP-Aux1 molecules recruited during the burst (Cohen’s *d* = 0.45) and in the lifetime (Cohen’s *d* = 0.57) of clathrin coated structures, determined in 1272 traces from 14 wild-type (WT) cells and in 794 traces from 14 knockout (GAK-KO) cells. (**b**) Effect of Aux1 knockdown by shRNA on recruitment of GAK to endocytic clathrin coated vesicles, in cells gene-edited for EGFP-GAK^+/+^ and CLTA-TagRFP^+/+^. Histogram and cumulative distributions for the number of EGFP-Aux1 molecules recruited during the burst (Cohen’s *d* = 0.73) and the lifetimes of the clathrin coated structures (Cohen’s *d* = 0.29), determined in 1498 traces from 15 control cells and in 1793 traces from 14 knockdown (Aux1-KD) cells. (**c**) Effect of GAK knockout and Aux1 knockdown by siRNA on recruitment of Aux1 to endocytic clathrin coated vesicles, in cells gene-edited for EGFP-Aux1^+/+^ and CLTA-TagRFP^+/+^ and knockout for GAK. Histogram distributions for the number of EGFP-Aux1 molecules recruited during the burst of clathrin coated structures, determined in 1794 traces from 20 control cells and in 465 traces from 47 knockdown (Aux1-KD) cells.

Combined depletion of GAK and Aux1 in gene-edited cells led to substantial loss of endocytic coated vesicles (Supplementary Fig. 4k) and inhibition of transferrin uptake (Supplementary Fig. 4l), as expected from published knockdown experiments (Hirst et al., 2008). The remaining clathrin coated structures recruited small bursts of 1-2 molecules of EGFP-Aux1 (Fig. 5c) suggesting that very few auxilins can recruit enough Hsc70 for uncoating.

### Uncoating dynamics mediated by auxilins lacking their PTEN-like domain

The experimental results in Fig. 4 show that the PTEN-like domains of Aux1 and GAK determine timing and amplitude of recruitment to coated vesicles. A truncated Aux1 that retains just the clathrin-binding and J domains (ΔPTEN Aux1) can nonetheless direct uncoating *in vitro* (Bocking et al., 2011), and ectopic expression of a GAK transgene encoding only the clathrin-binding and J-domains (ΔPTEN GAK) reverses the lethality of a conditional GAK knockout in liver or brain of mice and restores clathrin traffic in embryo fibroblasts derived from those mice (Park et al., 2015). We therefore compared the dynamics of recruitment and the compartment specificity of wild-type and ΔPTEN auxilins in our SUM159 cells. Ectopic expression of various ΔPTEN variants of Aux1 or GAK, in cells devoid of GAK and depleted of Aux1, rescued transferrin uptake (Supplementary Fig. 5a). In these cells, ectopically expressed ΔPTEN EGFP-Aux1 or ΔPTEN EGFP-GAK also exhibited a burst of recruitment, with amplitude of 1-5 EGFPs, just after completion of coat assembly (Supplementary Fig. 5b,c). We chose for analysis cells with levels of ectopic expression comparable to those in gene-edited cells expressing the fluorescent auxilin under control of the endogenous promoter. Association with coated vesicles adequate to drive uncoating thus did not require PTEN-like domain recognition of phosphoinositides. The mean burst amplitude was comparable to the lowest amplitudes seen with full-length Aux1 or GAK, confirming that Hsc70 recruitment by relatively few auxilins can drive uncoating.

The ΔPTEN EGFP-Aux1 in the experiments just described lacked the PTEN lipid sensor, yet it appeared only in fully assembled coated vesicles and not in coated pits. In our previous paper (He et al., 2017), we showed that probes with the Aux1 clathrin-binding domain and a sensor for PtdIns(4,5)P_2_ or PtdIns(4)P (phophoinositides present in the plasma membrane) do appear in coated pits. Thus, during coated pit formation, recognition of the clathrin lattice appears to be necessary but not sufficient for auxilin recruitment; additional lipid-headgroup affinity is also required. After budding, however, clathrin coated vesicles can recruit Aux1 or GAK lacking any PTEN-like domain. The average recruitment levels for these species in our experiments was substantially lower than for the corresponding wild-type proteins, but it was nevertheless sufficient to elicit uncoating.

We carried out similar experiments to show that GAK lacking its PTEN-like domain can rescue the AP1 coated-vesicle dispersal phenotype seen previously in HeLa cells depleted of full-length GAK (Kametaka et al., 2007; Lee et al., 2005) (Supplementary Fig. 5d,e). Low-level ectopic expression of ΔPTEN GAK in cells gene-edited to eliminate endogenous GAK led to recovery of the perinuclear distribution of AP1 seen in wild-type cells (Fig. 2d and Supplementary Fig. 5e). A PTEN-less GAK thus appears to allow normal coated vesicle function in the secretory pathway.

### Phosphoinositide binding preferences of Aux1 and GAK

We sought further functional confirmation for the role of phosphoinositides in the uncoating reaction by using a previously established *in vitro* based uncoating assay designed to follow by single-object TIRF microscopy the ATP-, Hsc70 and auxilin dependent release of fluorescent clathrin from membrane-free clathrin/AP2 coats immobilized on a glass coverslip (Bocking et al., 2011). Binding of the Aux1 or GAK PTEN-like domains with phosphoinositides is relatively weak, and we turned to this assay when the conventional lipid-strip binding or vesicle floatation assays proved unreliable.

We modified as follows our previous assay, in which disassembly of clathrin/AP2 coats, with no enclosed liposome and loaded with saturating amounts of PTEN-less Aux1 (ΔPTEN-Aux1), was induced by addition of physiological amounts of ATP and Hsc70. First, we induced uncoating by simultaneous addition of recombinant ΔPTEN-Aux1, full length Aux1 or GAK (∼25 nM, an approximate physiological level) together with Hsc70 (1 μM) in the presence of 2 mM ATP. Second, we used synthetic clathrin/AP2 coated vesicles generated by assembling clathrin/AP2 coats onto pre-formed liposomes, ∼50-80 nm in diameter, doped with a peptidolipid bearing the YQRL endocytic motif (recognized by AP2) as well as with PtdIns(4,5)P_2_ (recognized by AP2) and either PtdIns(3)P (preferentially recognized by Aux1) or PtdIns(4)P (preferentially recognized by GAK) (Fig. 6a,b and Supplementary Fig. 6); a representative example of a time series is shown in Supplementary Video 4. We included trace amounts of one of two fluorescent lipid dyes in each type of liposome to identify it uniquely and to distinguish between synthetic clathrin/AP2 coated vesicles and clathrin/AP2 coats; we labeled clathrin with a third fluorescent dye (Fig. 6c). The relative amounts of clathrin associated with each coat, before and during the uncoating reaction were determined, as illustrated by the representative traces in Fig. 6d, from which we obtained the dwell time for the uncoating reaction (i.e., the interval between the first exposure to the uncoating mixture and initiation of clathrin release) (Fig. 6d) and the efficiency of uncoating (how much fluorescent clathrin was released from a given synthetic clathrin/AP2 coated vesicle during the 150 s duration of the time series) (Supplementary Fig. 6, Supplementary Video 4). Under these conditions, most clathrin/AP2 coats disassembled rapidly, in agreement with our earlier results from an *in vitro* single-object uncoating assay (Bocking et al., 2011; Bocking et al., 2018). In contrast, uncoating of the synthetic clathrin/AP2 coated vesicles was generally slower, and we could often detect partial release of the clathrin coat in what appeared to be steps (Fig. 6d). As expected for the control experiment carried out with the ΔPTEN-Aux1 fragment unable to recognize lipids but retaining the clathrin-, AP2- and Hsc70-binding domains, dwell time and uncoating efficiency were equivalent for the clathrin/AP2 coats and for the PtdIns(3)P- and PtdIns(4)P-containing synthetic clathrin/AP2 coated vesicles (Fig. 6e, left panel; Supplementary Fig. 6). Uncoating induced in the same assay by full length Aux1 initiated more rapidly for synthetic coated vesicles containing PtdIns(3)P (dwell time ∼5 s) than it did for synthetic coated vesicles containing PtdIns(4)P (dwell time ∼16 s) (Fig. 6e, middle panel), while uncoating induced by GAK initiated more rapidly for synthetic vesicles containing PtdIns(4)P (dwell time ∼20 s) than it did for those containing PtdIns(3)P (dwell time ∼29 s) (Fig. 6e, right panel). The observed dwell times, which we expect to depend on the kinetics of Aux1 or GAK recruitment, thus varied with lipid composition (the only variable in the comparisons); the uncoating efficiency, which should depend only on the ultimate arrival of Hsc70, did not (Supplementary Fig. 6). We conclude that the PTEN-like domains of Aux1 and GAK have phosphoinositide preferences for PtdIns(3)P and PtdIns(4)P, respectively.

**Figure 6.**
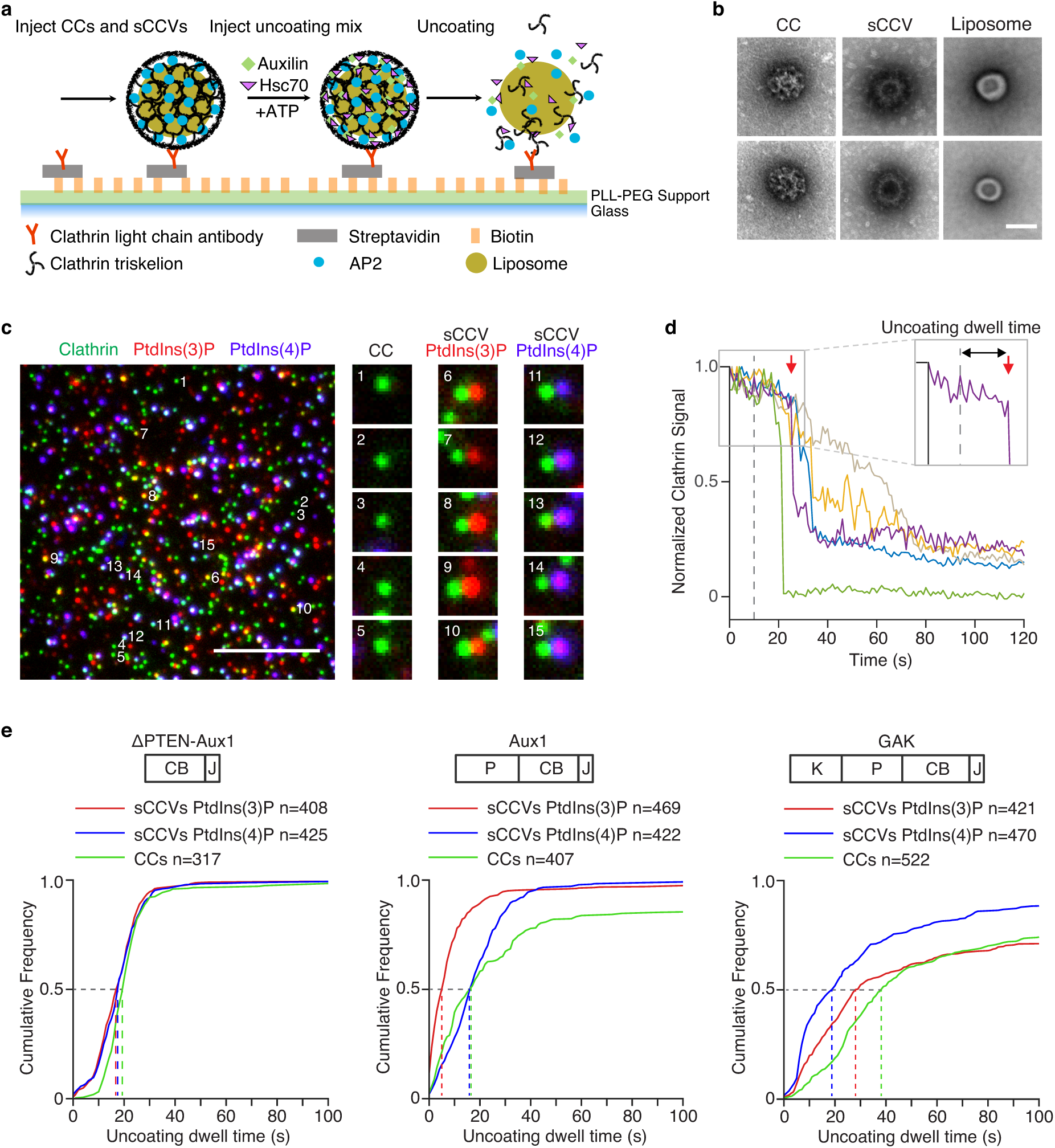
Single-object *in vitro* uncoating assay. (**a**) Schematic representation of the single-object uncoating assay. The intensities of fluorescence from labeled clathrin and of lipid dyes incorporated into liposomes surrounded by clathrin/AP2 coats were monitored by TIRF microscopy. Synthetic clathrin/AP2 coated vesicles (shown in the figure) and clathrin/AP2 coats were captured with a biotinylated monoclonal antibody specific for clathrin light chain on the surface of a PLL-PEG-Biotin-Streptavidin modified glass coverslip in a microfluidic chip. (**b**) Representative transmission electron microscopy images of negatively stained clathrin/AP2 coats (CC) (left), synthetic clathrin/AP2 coated vesicles (sCCV) (middle) or liposomes (right). Scale bar, 50 nm. (**c**) Representative TIRF image before initiation of the single-object uncoating assay. The snapshot combines images acquired in three different fluorescence channels (red and blue channels shifted right by 5 pixels) used to monitor the signal from coats and synthetic coated vesicles tagged with clathrin LCa-AF488 (green) and synthetic coated vesicles containing PtdIns(3)P- or PtdIns(4)P and DiI (red) or DiD (blue), respectively. Scale bar, 5 µm. (**d**) Representative uncoating profiles from single synthetic clathrin/AP2 coated vesicles. The plots show fluorescence intensity traces of the clathrin signal imaged at 1 s intervals starting 10 s prior to arrival of the uncoating mix (dashed vertical line at 0 s). The abrupt loss of signal in the green trace represents early release of the synthetic coated vesicle from the antibody on the glass surface. The enlarged boxed region (right corner) illustrates with a red arrow the onset of the uncoating reaction of the purple trace; the time it took to reach this point is the defined as the uncoating dwell time. (**e**) Cumulative distributions of uncoating dwell times of clathrin/AP2 and synthetic clathrin/AP2 coated vesicles containing PtdIns(4,5)P_2_ with either PtdIns(3)P or PtdIns(4)P obtained upon incubation with ΔPTEN-Aux1, full length Aux1, or full length GAK (P: PTEN-like domain; CB: clathrin-binding domain; J: J domain; K: kinase domain). The uncoating dwell times corresponding to 50% of the distributions are indicated (dashed lines) are from 3-10 independent experiments.

## DISCUSSION

The experiments described here have yielded several unexpected findings. One concerns the absence of detectable Aux1 or GAK during the assembly phase of the coated pits. Past experiments done by ectopic expression of Aux1 or GAK, often resulted in recruitment of Aux1 or GAK during the assembly phase; some have interpreted this recruitment as a way to explain the partial exchange of clathrin observed during coat assembly concluding by inference that this is an Hsc70 mediated process. Here, we have combined single-molecule live-cell imaging sensitivity with physiological expression of fluorescently tagged Aux1 and GAK to show absence of Aux1 and GAK during coated pit assembly (Supplementary Fig. 2c-f). These observations resolve a long-standing discussion by demonstrating that Aux1 and/or GAK cannot explain the exchange of clathrin during pit formation (and by inference that Hsc70 likewise has no role).

The second discovery concerned determination of the stoichiometry by which Aux1 and GAK are recruited to the clathrin/AP2 coat during uncoating. Understanding the extent of this recruitment is fundamental to understanding the mechanism of the uncoating process, and because Hsc70 is a “disassemblase” for many important cellular processes. We found that relatively few auxilins were sufficient for functional uncoating. Calibrated measurements showed that peak occupancy by 3-4 molecules of Aux1 or GAK yielded complete uncoating and that only rarely was the Aux1 or GAK occupancy higher. Uncoating initiated with even fewer auxilins, and the maximum occupancy generally occurred after uncoating had begun, coinciding in average “cohort” traces with roughly 50% loss of clathrin (Fig. 1d,g). Moreover, when we eliminated GAK by gene editing and depleted Aux1 by RNAi, we found, in the cells with slightly incomplete knockdown, a maximal occupancy at any one time of just 1-2 Aux1 molecules per clathrin-coated vesicle, which nonetheless appeared to uncoat with normal kinetics.

High-affinity engagement of substrate by Hsc70 requires both a bound J-domain and ATP hydrolysis. Activated Hsc70 can therefore associate only at a vertex adjacent to its activating auxilin. The structure of Hsc70-bound coats suggests a vertex can accommodate no more than one Hsc70 (Xing et al., 2010). Auxilin occupancy is therefore a first-level estimate of the number of Hsc70s required for functional uncoating *in vivo*. If the time for Hsc70 recruitment, clathrin binding, ATP hydrolysis, and release from the J-domain is shorter than the uncoating time, however, the peak steady-state level of auxilin will sometimes underestimate the total number of auxilins that have participated. That is, dissociation of auxilin from the neighborhood of one vertex, having delivered its Hsc70, and vicinal re-binding (or binding of a different auxilin) at another vertex could result in an apparent steady-state occupancy of only one auxilin but delivery of Hsc70s to two distinct vertices. Some individual traces of the few events we could find in GAK knockout cells with essentially complete Aux1 knockdown suggested that two or three single Aux1 molecules might have arrived independently during the course of uncoating. Putting together data from the various regimes we have examined, we estimate that recruitment of no more than 3-5 auxilins, and of probably fewer than 10 Hsc70s, is enough to dismantle coated vesicles in the size range (∼60 clathrin trimers) present in the cells we have used. Under normal conditions of Aux1 or GAK expression, additional Hsc70s might participate. A recent model derived from ensemble *in vitro* uncoating experiments concluded that stoichiometric amounts of Aux1/GAK with respect to clathrin were recruited in order to mediate a proposed sequential capture of up to three Hsc70 molecules to each triskelion (Rothnie et al., 2011). We have now ruled out this model by the *in vivo* counting of the number of molecules of Aux1/GAK recruited to a coated vesicle during uncoating. Our data clearly show that the recruitment is sub-stoichiometric -- indeed, ∼30% of all uncoating events occur with less than eight-recruited Aux1/GAK. Our *in vivo* data instead agree with earlier biochemical studies suggesting a ‘catalytic’ role for Aux1 and GAK during uncoating (Ma et al., 2002).

In previous *in vitro* single-molecule uncoating experiments, which were carried out by saturating the coats with auxilin and then adding Hsc70, we found from kinetic modeling that uncoating occurred precipitously when the Hsc70 level had reached an occupancy of about one for every two vertices (Bocking et al., 2011). Since auxilin itself stabilizes coats (Ahle and Ungewickell, 1990), this level is likely to be substantially greater than the Hsc70 occupancy needed to uncoat with limiting auxilin present. To address this question directly, we modified our single-molecule uncoating reaction in two ways: first, by inducing uncoating with a mixture of 1 μM Hsc70 and 25 nM Aux1 or GAK (roughly physiological concentrations (Kulak et al., 2014), and second, by including as substrates synthetic clathrin/AP2 coated vesicles containing PtdIns(3)P or PtdIns(4)P. This assay is a sensitive functional test of how the kinetics of uncoating depends on lipid composition. We found that Aux1 initiated uncoating of vesicles containing PtdIns(3)P more rapidly than uncoating of vesicles containing PtdIns(4)P; GAK had the opposite preference. The onset of uncoating mediated by Aux1, GAK or ΔPTEN-Aux1 was the same when using as substrate clathrin/AP2 coats lacking any encapsulated liposome (Fig. 6). This functional assay was proved substantially more robust for the relatively low-affinity PTEN-like domains than did liposome or lipid-strip binding assays.

During uncoating, loss of clathrin and loss of AP2 follow each other closely. AP2 adheres to the plasma membrane by virtue of its affinity of PtdIns(4,5)P_2_, which is hydrolyzed (to PtdIns(4)P) by synaptojanin (in coated pits) and OCRL (in coated vesicles) (Chang-Ileto et al., 2011; He et al., 2017; Nandez et al., 2014). But only after pinching off of the vesicle does cessation of rapid exchange allow the PtdIns(4,5)P_2_ concentration to fall and the PtdIns(4)P concentration to rise (He et al., 2017), reducing AP2 affinity for the vesicular membrane. Because AP2 also stabilizes clathrin association with the vesicle (and hence with other clathrins) (Kirchhausen et al., 2014), PtdIns(4,5)P_2_ depletion after pinching may accelerate uncoating under conditions of limiting auxilin.

Auxilin binding requires contributions from three different clathrin trimers organized in the lattice of a coat (Fotin et al., 2004a). Our results, together with our previously published work (He et al., 2017), show that coincident recognition of this local structure and of the cognate lipid determines the timing of normal Aux1 and GAK recruitment. Nonetheless, deletion of the PTEN-like domain did not fully disable auxilin association, which occurred as a low-amplitude burst following completion of coat assembly, with no evidence of premature association. Interactions other than with PtdIns(3)P (for Aux1) and PtdIns(4)P (for GAK) must therefore contribute to the observed dependence on vesicle closure. One possibility could be a slightly different structure (e.g., a “tighter” one, due to closure) of the clathrin lattice associated with coated vesicles resulting in enhanced accessibility of the Aux1/GAK binding regions in the clathrin terminal domain or in AP2 (Scheele et al., 2001). Our current data, however, offer no definitive evidence for the source of this redundancy.

Finally, concerning the dynamics of coated pit / coated vesicle formation, we have shown a straightforward way to distinguish abortive coated pits (i.e., those that fail to form coated vesicles) from coated pits that mature and become coated vesicles. The best currently available method relies on a detailed analysis of the distribution of coated-pit lifetimes (Aguet et al., 2013). Because we have now shown that Aux1/GAK are recruited only to coated vesicles and not to assembling coated pits, we can simply segregate clathrin or AP2 traces by whether or not they end with an Aux1/GAK burst. This simple assignment, similar to the recruitment of dynamin (Aguet et al., 2013; Ehrlich et al., 2004), provides a robust way to distinguish between dissociation of the lattices of abortive pits and disassembly of the lattices of coated vesicles and hence between an abortive pit (whatever its lifetime) and one that proceeds to pinch off as a coated vesicle. We further note that the outcome of this analysis has shown in a simple way that the distinction between abortive and non-abortive events is a meaningful one.

## METHODS

### Antibodies

The antibody against Aux1/GAK was a kind gift from Sanja Sever (Massachusetts General Hospital, Harvard Medical School) (Newmyer et al., 2003). The antibody against GAK (M057-3) was purchased from MBL International Corp.

### Cell culture

The mostly diploid SUM159 human breast carcinoma cells (Forozan et al., 1999) kindly provided by J. Brugge (Harvard Medical School) were routinely verified to be mycoplasma free using a PCR-based assay. SUM159 cells were grown at 37°C and 5% CO_2_ in DMEM/F-12/GlutaMAX (GIBCO, Langley, OK), supplemented with 5% fetal bovine serum (FBS, Atlanta Biologicals, Lawrenceville, GA), 100 U/ml penicillin and streptomycin (VWR International, Philadelphia, PA), 1 µg/ml hydrocortisone (Sigma-Aldrich, St. Louis, MO), 5 µg/ml insulin (Sigma-Aldrich, St. Louis, MO), and 10 mM HEPES (Mediatech, Manassas, VA), pH 7.4.

### Plasmids, transfection and ectopic expression

The DNA sequences encoding the full-length bovine Aux1 (910 residues, NM_174836.2), or full-length human GAK (1311 residues, NM_005255.3) were amplified by PCR from full-length cDNA clones (Massol et al., 2006) and inserted into pEGFP-C1 or mCherry-C1 to generate the plasmids EGFP-Aux1, mCherry-Aux1, EGFP-GAK and mCherry-GAK. The kinase domain (residues 1-347 of GAK), PTEN-like domain (residues 1-419 of Aux1, residues 360-766 of GAK), clathrin-binding domain (residues 420-814 of Aux1; residues 767-1222 of GAK) and J domain (residues 815-910 of Aux1; residues 1223-1311 of GAK) were amplified by PCR from the full-length cDNA clones (Massol et al., 2006) and inserted into pEGFP-C1 to generate the EGFP-fused Aux1 or GAK truncations. These DNA segments were also fused by overlap PCR and inserted into pEGFP-C1 to generate the EGFP-Aux1/GAK chimera. All constructs used the linker (5’-GGAGGATCCGGTGGATCTGGAGGTTCTGGTGGTTCTGGTGGTTCC-3’) placed between the DNA fragments and EGFP or mCherry.

Transfections were performed using TransfeX Transfection Reagent (ATCC, Manassas, VA) according to the manufacturer’s instructions and cells with relatively low levels of protein expression were subjected to live cell imaging 16-20 hours after transfection.

### Genome editing of SUM159 cells to express EGFP-Aux1^+/+^, EGFP-GAK^+/+^, TagRFP-GAK^+/+^, or AP1-TagRFP^+/+^ using the CRISPR/Cas9 approach

SUM159 cells were gene-edited to incorporate EGFP or TagRFP to the N-terminus of Aux1 or GAK, or the C-terminus of AP1 sigma-1 subunit using the CRISPR/Cas9 approach (Ran et al., 2013). The target sequences overlapping the start codon ATG (underlined) at the genomic locus recognized by the single-guide RNA (sgRNA) are 5’-<u>ATG</u>AAAGATTCTGAAAATAA-3’ for *DNAJC6* (encoding Aux1) and 5’-CGCC<u>ATG</u>TCGCTGCTGCAGT-3’ for *GAK*. The target sequence overlapping the stop codon TAG (underlined) at the genomic locus recognized by the sgRNA is GGTTTGGCA<u>TAG</u>CCCCTGCT for *AP1S1*. The sgRNA containing the targeting sequence was delivered as PCR amplicons containing a PCR-amplified U6-driven sgRNA expression cassette (Ran et al., 2013).

The donor constructs EGFP-Aux1 and EGFP-GAK used as templates for homologous recombination to repair the Cas9-induced double-strand DNA breaks were generated by cloning into the pCR8/GW TOPO vector with two ∼800-nucleotide fragments of human genomic DNA upstream and downstream of the start codon of *DNAJC6* or *GAK* and the open reading frame of EGFP by TA ligation cloning. The upstream and downstream genomic fragments were generated by PCR amplification reactions from the genomic DNA extracted from SUM159 cells using the QiaAmp DNA mini kit (Qiagen). The open reading frame encoding EGFP together with a flexible linker (5’-GGAGGTTCTGGTGGTTCTGGTGGTTCC-3’) was obtained by PCR from an EGFP expression vector.

The donor constructs TagRFP-GAK and AP1-TagRFP were generated by cloning into the pUC19 vector with two ∼800-nucleotide fragments of human genomic DNA upstream and downstream of the start codon of *GAK* or the stop codon of *AP1S1* and the open reading frame of TagRFP using the Gibson Assembly Master Mix (New England BioLabs). The open reading frames encoding TagRFP together with a flexible linker (5’-GGAGGATCCGGTGGATCTGGAGGTTCT-3’) were obtained by PCR from a TagRFP expression vector.

Clonal cell lines expressing EGFP-Aux1^+/+^, EGFP-GAK^+/+^, TagRFP-GAK^+/+^, or AP1-TagRFP^+/+^ was generated as described (He et al., 2017). In brief, SUM159 were transfected with 800 ng each of the donor plasmid, the plasmid coding for the Streptococcus pyogenes Cas9 and the free PCR product using Lipofectamin2000 (Invitrogen) according to the manufacturer’s instruction. Then the cells expressing EGFP or TagRFP chimeras were enriched by fluorescence-activated cell sorting (FACS) using a FACSAria II instrument (BD Biosciences), and further subjected to single cell sorting to select monoclonal cell populations. The cells with successful incorporation in the genomic locus of EGFP or TagRFP were screened by PCR using GoTaq Polymerase (Promega).

### Knockout of GAK using the CRISPR/Cas9 approach

Knockout of GAK was performed using the CRISPR/Cas9 approach exactly as described before (He et al., 2017) except that the target sequence for GAK overlapping the start codon ATG (underlined) is 5’-CGCC<u>ATG</u>TCGCTGCTGCAGT-3’.

### mRNA depletion of Aux1 and GAK by shRNA or siRNA knockdown

Lentivirus shRNA expressing 5’-CACTTATGTTACCTCCAGAAT-3’ or 5’-GAAGATCTGTTGTCCAATCAA-3’ was used to knock down the expression of Aux1 or GAK (Broad Institute TRC library) as described before (He et al., 2017); 5’-CCTAAGGTTAAGTCGCCCTCG-3’ was used as control. Alternatively, siRNAs were used to knockdown Aux1 or GAK using Lipofectamine RNAiMAX (Invitrogen). siGENOME SMARTpool (a mixture of four siRNAs) was used to knockdown GAK (M-005005-02-0005; Dharmacon); a single siGENOME siRNA was used to knockdown Aux1 (D-009885-02-0010; Dharmacon). A non-targeting siRNA (D-001210-03-05; Dharmacon) was used as a control. Knockdown of Aux1 or GAK by siRNA was achieved by two sequential transfections, the first one in cells after overnight plating and the second two days later, followed by analysis during the fourth day.

### Transferrin uptake by flow cytometry

Transferrin uptake by a flow cytometry–based assay was done as described (Cocucci et al., 2014).

### Western blot analysis

Western blot analysis was performed as described (Cocucci et al., 2012) using the anti-GAK antibody (1:500) or anti-Aux1/GAK antibody (1:500) diluted in Tris-buffered saline with Tween 20 containing 3% BSA.

### TIRF microscopy and spinning disk confocal microscopy: live-cell imaging and image analysis

The TIRF and spinning disk confocal microscopy experiments were as described (Cocucci et al., 2012). The single EGFP or TagRFP molecule calibration was carried out as described before (Cocucci et al., 2012; Cocucci et al., 2014). Recombinant EGFP made in *E. Coli* was used to determine the fluorescence intensity of a single EGFP molecule. The cytosol from gene-edited SUM159 cells expressing AP2-TagRFP^+/+^ was used to determine the fluorescence intensity of a single TagRFP molecule.

The detection and tracking of all fluorescent traces was carried out using the cmeAnalysis software package (Aguet et al., 2013). For the automated detection, the minimum and maximum tracking search radius were 1 and 3 pixels, and the maximum gap length in a trajectory was 2 frames (He et al., 2017). Detection of an independent event was verified by establishing absence of significant signal during brief intervals preceding and following the first and last detected signals. The intensity-lifetime cohorts were generated as described (Aguet et al., 2013). Detection and tracking of clathrin-coated structures and the associated Aux1 or GAK were carried out with clathrin or AP2 as the “master” and Aux1/GAK as the “slave” channel. The valid clathrin or AP2 traces with significant Aux1/GAK signal in the slave channel were selected automatically and verified manually. The amplitude of the 2-D Gaussian PSF fitting for the detected EGFP-Aux1 or EGFP-GAK was used to estimate the number of EGFP-Aux1 or EGFP-GAK molecules calibrated by the intensity of single EGFP. Detection and tracking of events in cells expressing only fluorescently tagged Aux1, GAK or lipid sensors were carried out with the following combinations of master and slave channels: PtdIns(4)P sensor/PtdIns(3)P sensor, PtdIns(3)P sensor/Aux1, PtdIns(4)P sensor/Aux1, PtdIns(3)P sensor/GAK, PtdIns(4)P sensor/GAK (Fig. 3a,c,d), EGFP-Aux1/TagRFP-GAK and mCherry-Aux1/EGFP-Aux1 (Fig. 3b,e), respectively. The validity of traces was verified manually. The frame associated with the maximum fluorescence intensities for these traces and the corresponding interval for each pair was determined automatically.

### Lattice light-sheet microscopy: live-cell imaging and image analysis

To show the subcellular localization of Aux1 and GAK in the whole cell volume, the gene-edited EGFP-Aux1^+/+^ and CLTA-TagRFP^+/+^ cells, the gene-edited EGFP-GAK^+/+^ and CLTA-TagRFP^+/+^ cells, and the EGFP-GAK^+/+^ and AP1-TagRFP cells were imaged using lattice light-sheet microscopy with a dithered square lattice light-sheet (Aguet et al., 2016; Chen et al., 2014). The cells were plated on 5 mm coverslips (Bellco Glass, Vineland, NJ) for at least 4 hours prior to imaging, and were imaged in FluoroBrite™ DMEM media (Thermo Fisher Scientific, Rockford, IL) containing 5% FBS and 20 mM HEPES at 37°C. The cells were sequentially excited with a 488 nm laser (300 mW) and a 560 nm laser (10-50 mW) for ∼100 ms for each channel using a 0.35 inner and 0.4 outer numerical aperture excitation annulus. The 3D volumes of the whole cells were recorded by scanning the sample at 250 nm step sizes in the s-axis (corresponding to ∼131 nm along the z-axis), thereby capturing a volume of ∼50 µm x 50 µm x 75 µm (512 x 512 x 300 pixels).

To track the dynamic recruitment of Aux1 or GAK in 3D, the gene-edited EGFP-Aux1^+/+^ and AP2-TagRFP^+/+^ cells, and the gene-edited EGFP-GAK^+/+^ and AP2-TagRFP^+/+^ cells, and the EGFP-GAK^+/+^ and AP1-TagRFP cells were imaged using lattice light-sheet microscopy. The cells were excited with a 488 nm laser for ∼50 ms using a 0.505 inner and 0.6 outer numerical aperture excitation annulus. The 3D volumes of the imaged cells were recorded by scanning the sample every ∼2.1 s for 187 s at 500 nm step sizes in the s-axis (corresponding to ∼261 nm along the z-axis), thereby capturing a volume of ∼50 µm x 50 µm x 15 µm (512 x 512 x 40 pixels).

### *In vitro* single-object uncoating

#### Protein production

The procedures to generate recombinant clathrin heavy chain produced in Sf9 cells, light chain produced in *E. coli* were as described (Bocking et al., 2011; Bocking et al., 2018). Labeling of light chain labeled with Alexa Fluor 488 and incorporation into recombinant clathrin triskelions were as described (Bocking et al., 2011). Recombinant Hsc70 and ΔPTEN-Aux1 were produced in *E. coli* and prepared as described (Rapoport et al., 2008).

The DNA sequences encoding full-length bovine Aux1 and full-length human GAK were flanked at the N-terminal (for Aux1) or C-terminal (for GAK) by 6x-His tags upon insertion into the pFastBac vector. Proteins were produced intracellularly in Sf9 cells following the Bac-to-Bac protocol (ThermoFisher Scientific). Cells were lysed by sonication or using a ball bearing bore homogenizer. Lysates were ultracentrifuged, and the supernatant was applied to nickel-NTA resin. Proteins were eluted with imidazole. Aux1 was further purified by gel filtration chromatography and concentrated using Millipore centrifugal devices.

#### Preparation of YQRL peptidolipids

The CKVTRRPKASDYQRLNL peptidolipid was prepared by adapting a previously described procedure (Bocking et al., 2018; Kelly et al., 2014). Briefly, a mixture of 20 mg/ml of YQRL containing peptide (prepared in 20 mM HEPES buffer pH 7.4), DMSO and maleimide-DOPE (1:1:2 v/v mixture respectively) was vortexed at 1000 rpm for 2 hours. The coupling reaction was quenched using 10 mM β-mercaptoethanol for 30 min. The YQRL peptidolipid was extracted by adding cholorform, methanol, and water (4:3:2.25 v/v mixture) followed by centrifugation at 1000 rpm for 5 min. The organic phase containing the peptidolipid was dried under argon and stored in a sealed argon atmosphere-containing vial. The films were re-suspended in chloroform and methanol mixture (2:1) at 2 mg/ml prior to use for liposome lipid film preparation.

#### Liposome preparation

All lipids (Avanti Polar Lipids, Alabaster, AL) were mixed in 20:9:1 chloroform:methanol:water and dried to prepare composition specific lipid films. Prior to formation of the synthetic clathrin-coated vesicles, the lipid film aliquots were hydrated in coated vesicle formation buffer (20 mM MES Hydrate pH 6.5, 100 mM NaCl, 2 mM EDTA, 0.4 mM DTT) to 300 µM final concentration and liposomes extruded with a 50 nm diameter pore filter.

#### In vitro reconstitution of synthetic clathrin-coated vesicles

The following procedure, used to generate synthetic clathrin/AP2 coated vesicles (sCCVs) (Bocking et al., 2018), was based on the co-assembly of a clathrin and AP2 coat surrounding liposomes: a solution containing recombinant clathrin heavy chain and fluorescently labeled light chain (1:3 mol/mol ratio) and AP2 (3:1 w/w clathrin:AP2) (100 ug of clathrin heavy chain) was added to 15 ul of extruded liposomes (300 umol lipid /300 µl) made of 86.9% DOPC, 5% PtdIns(4,5)P_2_, 5% PtdIns(4)P or PtdIns(3)P, 3% YQRL DOPE peptidolipid, 0.1% DiI or DiD lipid dye and dialyzed overnight at 4 °C against coated vesicle formation buffer (80mM MES pH 6.5, 20mM NaCl, 2mM EDTA, 0.4mM DTT) using a Slide-A-Lyzer mini dialysis device (10K molecular weight cutoff, Thermo Fisher Scientific) followed by for an additional 4 hours of dialysis using fresh coated vesicle formation buffer. Large aggregates were removed by centrifugation using a bench top Eppendorf centrifuge at 4 °C for 10 min at maximum speed. The supernatant was then transferred to a fresh tube and centrifuged at high speed in a TLA-100.4 centrifuge (Beckman) at 70000 rpm for 30 min. The sCCV-containing pellet was re-suspended in coated vesicle formation buffer and centrifuged a second time at 70000 rpm for 30 min. The pellet was re-suspended using coated vesicle formation buffer (100 µl of buffer per 100 µg of clathrin heavy chain) and stored at 4 °C for up to one week.

#### Transmission electron microscopy

sCCVs were adsorbed for 60 s onto freshly glowed-discharged carbon coated electron microscope grids, washed with a few drops of Milli-Q water, stained for 30 s with a few drops 1.2% uranyl acetate and blot dried. The samples were imaged on a Tencai G^2^ Spirit BioTWIN (FEI, Hillsboro, OR) at 23000-49000 x magnification.

#### Microfluidic uncoating chamber preparation

Microfluidic chips (Bocking et al., 2011; Bocking et al., 2018) with PDMS plasma bonded to glass coverslips suited for TIRF microscopy were employed to efficiently deliver reagents to uncoat immobilized sCCVs. Glass coverslips (#1.5) were cleaned for a total of 20 min by sequential incubation in the following solvents: Toluene, Dichloromethane, Ethanol, Ethanol:Hydochloric Acid (1:1 v/v), and Milli-Q water. The clean coverslips were oxygen plasma treated and bonded to PDMS channels. These chips were immediately incubated with 1 mg/ml biotinylated PLL-PEG for 5 min, washed with Milli-Q water, and treated with streptavidin (20 µl of 1 mg/ml streptavidin dissolved in PBS added to 80 µl 20 mM Tris pH 7.5, 2 mM EDTA, 50 mM NaCl) for 5 min. The chips were functionalized with CVC.6 biotinylated antibody specific for clathrin light chain A as previously described (Bocking et al., 2011; Bocking et al., 2014).

#### In vitro uncoating of synthetic clathrin-coated vesicles

sCCVs were bound to the functionalized upper surface of the glass cover slip and those that failed to attach washed away by flowing uncoating buffer (20 mM imidazole pH 6.8, 100 mM KCl, 2 mM MgCl_2_, 5 mM protocatechuic acid, 50 nM protocatechuate-3,4-dioxygenase, 2 mM Trolox, 8 mM 4-Nitrobenzyl Alcohol). Disassembly of the clathrin/AP2 coats was caused with uncoating buffer supplemented with 1 µM Hsc70, 5 mM ATP, 10 mM MgCl_2_ and the appropriate auxilins (typically 25 nM of ΔPTEN-Aux1, full length Aux1 or full length GAK) flown through the chip at 20 µl/min.

The total internal fluorescence angle was set to the value that led to 80% of the maximal fluorescence signal observed for immobilized clathrin/AP2 coats and sCCVs before uncoating. The clathrin signal was monitored by exciting the Alexa Fluor 488 maleimide covalently linked to the clathrin light chain (Bocking et al., 2011). Time series starting 10 s prior to the uncoating mix and lasting 150 s were then recorded at an interval of 1s. Liposomes containing PtdIns(4)P- or PtdIns(3)P were independently labeled with DiI and DiD (Thermo Fisher Scientific) lipid dyes and detected by excitation at 561 nm and 640 nm, respectively. Signals from empty clathrin coats, PtdIns(3)P- and PtdIns(4)P-containing coated vesicles were classified using the 2D point source detector previously described (Aguet et al., 2013). The start of uncoating (the time point marking the onset of loss of the clathrin fluorescence signal) was manually curated for all traces included in the analysis.

### Statistical tests

Because the large size of the sample sizes, the Cohen’s *d* effect size (Cohen, 1988) was used to report the practical significance of the difference in the magnitude between the recruited EGFP-Aux1 or EGFP-GAK molecules and the lifetime of coated pits before and after the knockout or knockdown of GAK and Aux1. To compare the means from the cells with different treatments, the two-tailed *t*-test or one-way ANOVA was used as indicated in figure legends.

## ACKNOWLEDGMENTS

We thank S. Sever for the Aux1/GAK antibody gift and Justin R. Houser for maintaining the TIRF and spinning disc microscopes. We specially thank S.C. Harrison for discussions and editorial help. We also thank members of our laboratory for help and encouragement. E.S. was supported by grants from the National Natural Science Foundation (Grant No. 31270884) and by the Youth Innovation Promotion Association, Chinese Academy of Sciences. T.K. acknowledges support from the Janelia Visitor Program and Eric Betzig, Eric Marino, Tsung-Li Liu and Wesley R. Legant for help and advice in constructing and installing the lattice light-sheet microscope. Construction of the lattice light-sheet microscope was supported by grants from Biogen and Ionis Pharmaceuticals to T.K. The research was supported by NIH grant NIH R01 GM075252 to T.K.

## AUTHOR CONTRIBUTIONS

K.H., E.S. and T.K. designed experiments; K.H., E.S. and S.D. generated the gene-edited cell lines and collected the imaging data using TIRF and spinning disk confocal microscopies; K.H. E.S. and S.D analyzed the data collected using TIRF and spinning disk confocal microscopies; S.U. and W.S. collected the imaging data using the lattice light-sheet microscope; S.U. and K.H. analyzed the imaging data from the lattice light-sheet microscope; K.H. E.S. and S.D generated the constructs for ectopic expression of proteins; M.M., R.G. and E.S. generated the constructs for genome editing; S.U., K.B., B.C. I.R, and I.K. participated in the preparation of reagents and acquisition of data associated with the *in vitro* single-object uncoating experiments. K.H. and T.K. contributed to the final manuscript in consultation with the authors.

---

**Supplementary Figure 1.**
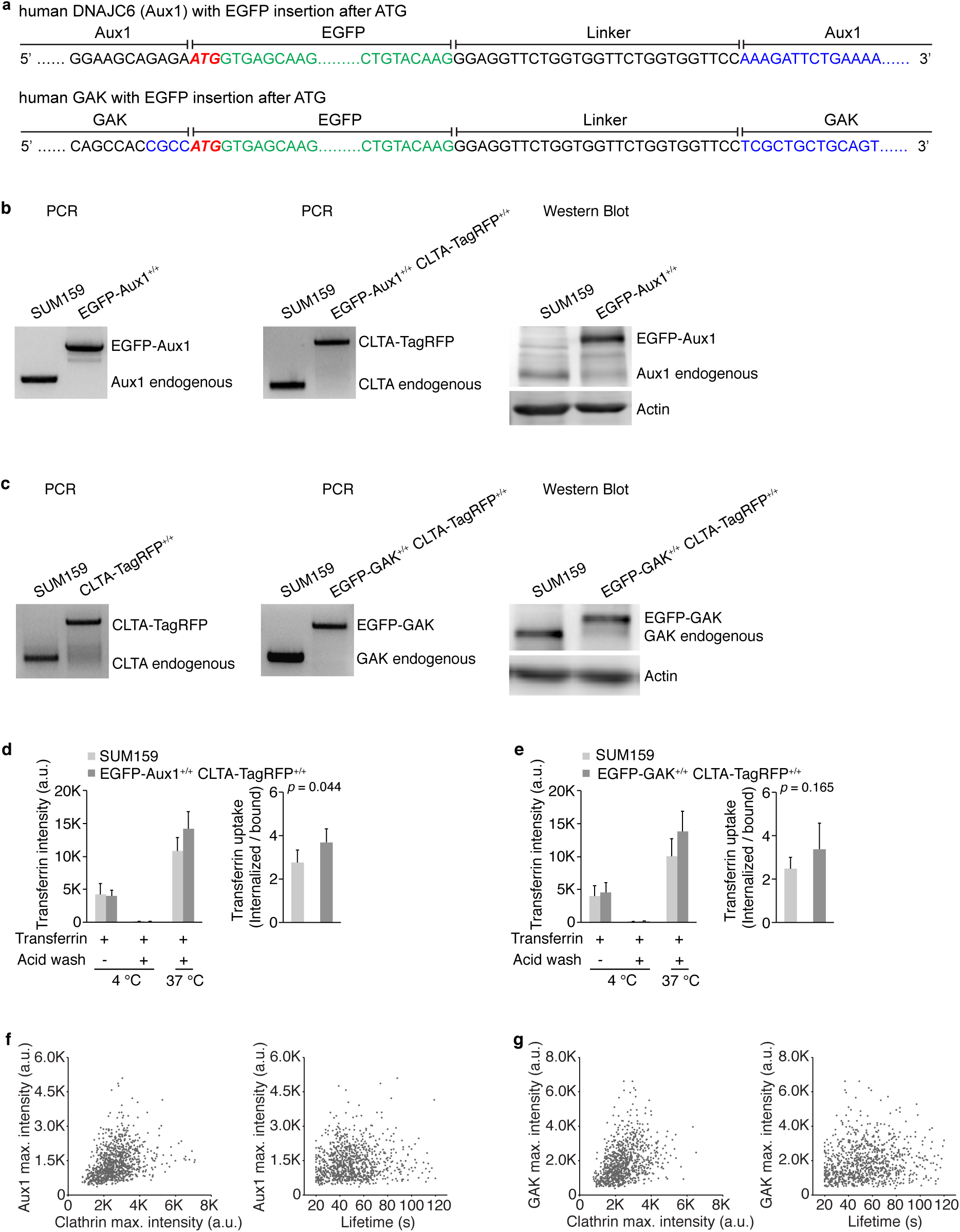
Gene-editing of SUM159 cells to express CLTA-TagRFP and EGFP-Aux1 or CLTA-TagRFP and EGFP-GAK. (**a**) CRISPR/Cas9 gene-editing strategy used to incorporate EGFP at the N-terminus of Aux1 or GAK. The resulting DNA sequences including the short linker between the C-terminus of EGFP and N-terminus of Aux1 or GAK are shown. (**b**) Genomic PCR analysis showing biallelic integration first of EGFP into the *DNAJC6* (Aux1) genomic locus to generate the clonal gene-edited cell line EGFP-Aux1^+/+^ (left panel) and then of TagRFP into the *CLTA* genomic locus of the same cells to generate the clonal double gene-edited cell line EGFP-Aux1^+/+^ CTLA-TagRFP^+/+^ (center panel). Right panel shows western-blot analysis of cell lysates from the EGFP-Aux1^+/+^ cells probed with antibodies for Aux1/GAK and actin. Although the genomic PCR shows biallelic integration of EGFP sequence into the Aux1 genomic locus, the western blot indicates expression of a small amount (∼15%) of untagged Aux1. The expression of EGFP-Aux1 in EGFP-Aux1^+/+^ cells was higher than endogenous Aux1 in the parental SUM159 cells; this up-regulation of EGFP-Aux1 expression, due either to single-cell cloning selection or to the genome editing. (**c**) Genomic PCR analysis showing biallelic integration first of TagRFP into the *CLTA* genomic locus to generate the clonal gene-edited cell line CLTA-TagRFP^+/+^ (left panel) and then of EGFP into the *GAK* genomic locus of the same cells to generate the clonal double gene-edited cell line EGFP-GAK^+/+^ and CTLA-TagRFP^+/+^ (center panel). Right panel shows western-blot analysis of cell lysates from the EGFP-GAK^+/+^ and CTLA-TagRFP^+/+^ cells probed with antibodies for GAK and actin. (**d**) Effect of expression of EGFP-Aux1 and CTLA-TagRFP on receptor-mediated uptake of transferrin. The histogram shows similar amounts of internalized Alexa Fluor 647-conjugated transferrin in parental and gene-edited EGFP-Aux1^+/+^ and CTLA-TagRFP^+/+^ cells probed by flow cytometry (n = 5 experiments, mean ± S.D., P value by two-tailed *t*-test). (**e**) Effect of expression of EGFP-GAK and CTLA-TagRFP on receptor-mediated uptake of transferrin (n = 5 experiments, mean ± S.D., P value by two-tailed *t*-test). (**f**) Scatter plots comparing maximum fluorescence intensities of EGFP-Aux1 and CLTA-TagRFP with each other (left panel; Pearson correlation coefficient *r* = 0.331) and maximum fluorescence intensity of EGFP-Aux1 with the lifetime of the endocytic coated structure in which it was found (right panel; Pearson correlation coefficient *r* = 0.115), from 938 traces in 8 cells. Data from bottom surfaces of double gene-edited EGFP-Aux1^+/+^ and CLTA-TagRFP^+/+^ cells imaged at 1 s intervals for 300 s by TIRF microscopy. (**g**) Scatter plots comparing maximum fluorescence intensities of EGFP-GAK and CLTA-TagRFP with each other (left panel; Pearson correlation coefficient *r* = 0.373) and maximum fluorescence intensity of EGFP-GAK with the lifetime of the endocytic coated structure in which it was found (right panel; Pearson correlation coefficient *r* = 0.153), from 900 traces in 8 cells. Data from bottom surfaces of double gene-edited EGFP-GAK^+/+^ and CLTA-TagRFP^+/+^ cells imaged at 1 s intervals for 300 s by TIRF microscopy.

**Supplementary Figure 2.**
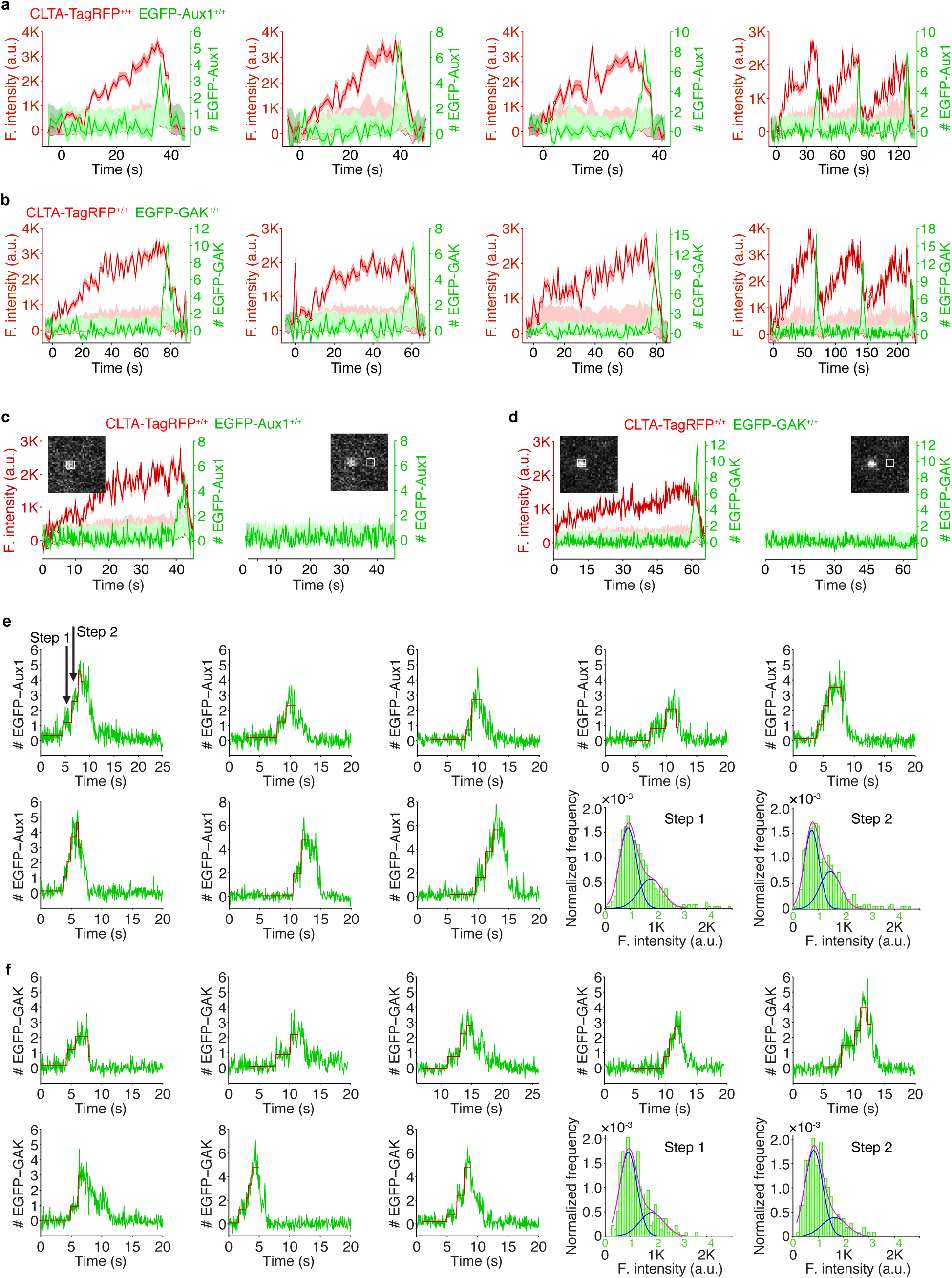
Recruitment of Aux1 and GAK to clathrin-coated vesicles in genome-edited cells. (**a**) Representative plots of single endocytic events (first 3 panels) and hotspots (last panel) showing fluorescence intensity traces for CLTA-TagRFP and EGFP-Aux1 (arbitrary units for CLTA; number of molecules for Aux1) imaged at 1 s intervals by TIRF microscopy. (**b**) Representative plots of single endocytic events (first 3 panels) and hotspots (last panel) showing fluorescence intensity traces for CLTA-TagRFP and EGFP-GAK imaged at 1 s intervals by TIRF microscopy. (**c**) EGFP-Aux1 recruitment was not detected while coated pits were assembling. Representative plots of a single endocytic event showing fluorescence intensity traces for CLTA-TagRFP and EGFP-Aux1 (left panel) imaged at 250 ms intervals by TIRF microscopy. The EGFP-Aux1 signal was detected and measured as indicated in the insert image. Right panel shows the fluorescence intensity fluctuations of the EGFP channel measured from the boxed area 12 pixels away from the detected EGFP-Aux1 burst signal. (**d**) EGFP-GAK recruitment was not detected while coated pits were assembling. (**e**) Stepwise recruitment of Aux1 to coated vesicles. Representative plots of EGPF-Aux1 burst-like recruitment (shown as number of molecules for Aux1) imaged at 62.5 ms intervals with TIRF microscopy; fit (red) obtained by applying a step-fitting function to estimate the average recruited molecules during the initiation phase of Aux1 burst-like recruitment. The last two panels show the histogram distributions (with Gaussian fitting) of EGFP-Aux1 molecules during the first step and second step of its recruitment. (**f**) Stepwise recruitment of GAK to coated vesicles.

**Supplementary Figure 3.**
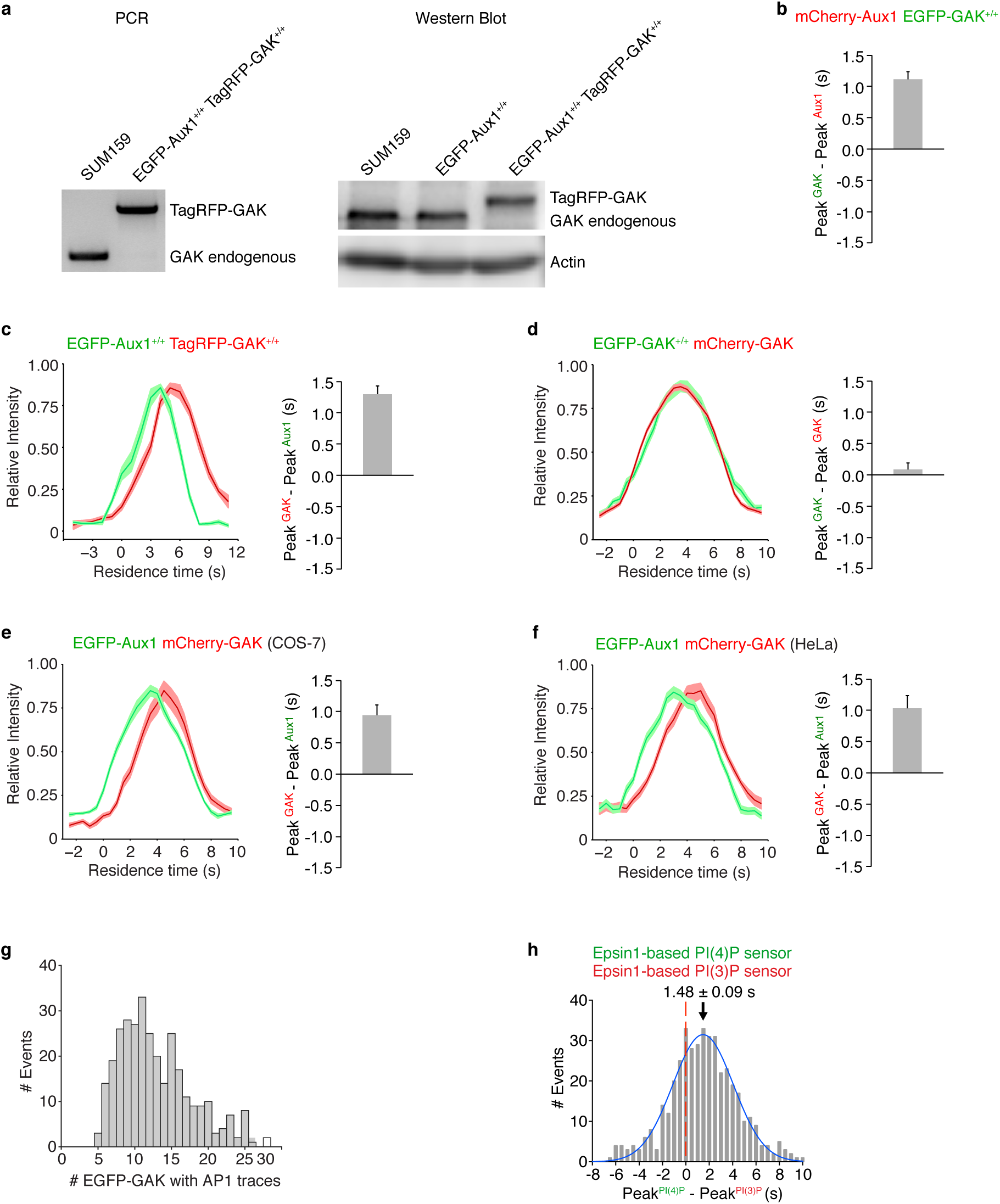
Sequential bursts of Aux1 and GAK during uncoating of clathrin-coated vesicles at the plasma membrane and recruitment of GAK to the intracellular clathrin-containing carriers. (**a**) The TagRFP sequence was inserted into the *GAK* genomic locus of the EGFP-Aux1^+/+^ cells to generate the double gene-edited cells EGFP-Aux1^+/+^ and TagRFP-GAK^+/+^, as confirmed by genomic PCR analysis (left panel) and western blot analysis probed with antibodies for GAK and actin (right panel). (**b**) Gene-edited EGFP-GAK^+/+^ cells transiently expressing mCherry-Aux1 were imaged at 0.5 s intervals for 60 s by TIRF microscopy. The average time interval between the peaks of intensity for EGFP-GAK and mCherry-Aux1 is shown (mean ± S.D., n = 8 cells). (**c**) Bottom surfaces of EGFP-Aux1^+/+^ and TagRFP-GAK^+/+^ cells were imaged at 1 s intervals for 120 s by TIRF microscopy. The left panel shows the averaged fluorescence intensity traces (mean ± S.E.) of both EGFP-Aux1 (green) and TagRFP-GAK (red) for the EGFP-Aux1 3–12 s cohort (1560 traces from 12 cells). The right panel shows the average time interval between the peaks of intensity for EGFP-Aux1 and TagRFP-GAK (mean ± S.D, n = 6 cells). (**d**) Gene-edited EGFP-GAK^+/+^ cells transiently expressing mCherry-GAK were imaged at 0.5 s intervals for 60 s by TIRF microscopy. The left panel shows the averaged fluorescence intensity traces (mean ± S.E.) of EGFP-GAK (green) and mCherry-GAK (red) from the EGFP-GAK 3–12 s cohort (2306 traces from 15 cells). The right panel shows the average interval between the peak intensities of EGFP-GAK and mCherry-GAK (mean ± S.D., n = 15 cells). (**e**) COS-7 cells transiently expressing EGFP-Aux1 and mCherry-GAK were imaged at 0.5 s intervals for 60 s by TIRF microscopy. The left panel shows the averaged fluorescence intensity traces (mean ± S.E.) of EGFP-Aux1 (green) and mCherry-GAK (red) from the EGFP-Aux1 3–12 s cohort (656 traces from 9 cells). The right panel shows the average interval between the peak intensities of EGFP-Aux1 and mCherry-GAK (mean ± S.D., n = 9 cells). (**f**) HeLa cells transiently expressing EGFP-Aux1 and mCherry-GAK were imaged at 0.5 s intervals for 60 s by TIRF microscopy. The left panel shows the averaged fluorescence intensity traces (mean ± S.E.) of EGFP-Aux1 (green) and mCherry-GAK (red) from the EGFP-Aux1 3–12 s cohort (595 traces from 11 cells). The right panel shows the average interval between the peak intensities of EGFP-Aux1 and mCherry-GAK (mean ± S.D., n = 11 cells). (**g**) Gene-edited EGFP-GAK^+/+^ cells stably expressing AP1-TagRFP were imaged in 3D by lattice light-sheet microscopy. Distribution of the maximum number of EGFP-GAK molecules recruited to individual AP1-coated carries (325 traces from 11 cells). (**h**) Bottom surfaces of cells transiently expressing Epsin1-based PtdIns(4)P sensor EGFP-P4M(DrrA)-Dlv2(508-736)-Epsin1(255-501) and PtdIns(3)P sensor mCherry-2xFYVE(Hrs)-Dlv2(508-736)-Epsin1(255-501) imaged by TIRF microscopy every 0.5 s for 100 s. Distribution (fit with a single Gaussian) for the interval between the peaks within single events showing that the Epsin1-based PtdIns(3)P sensor precedes the PtdIns(4)P sensor by 1.48 ± 0.09 s (mean ± S.E., 436 traces from 23 cells).

**Supplementary Figure 4.**
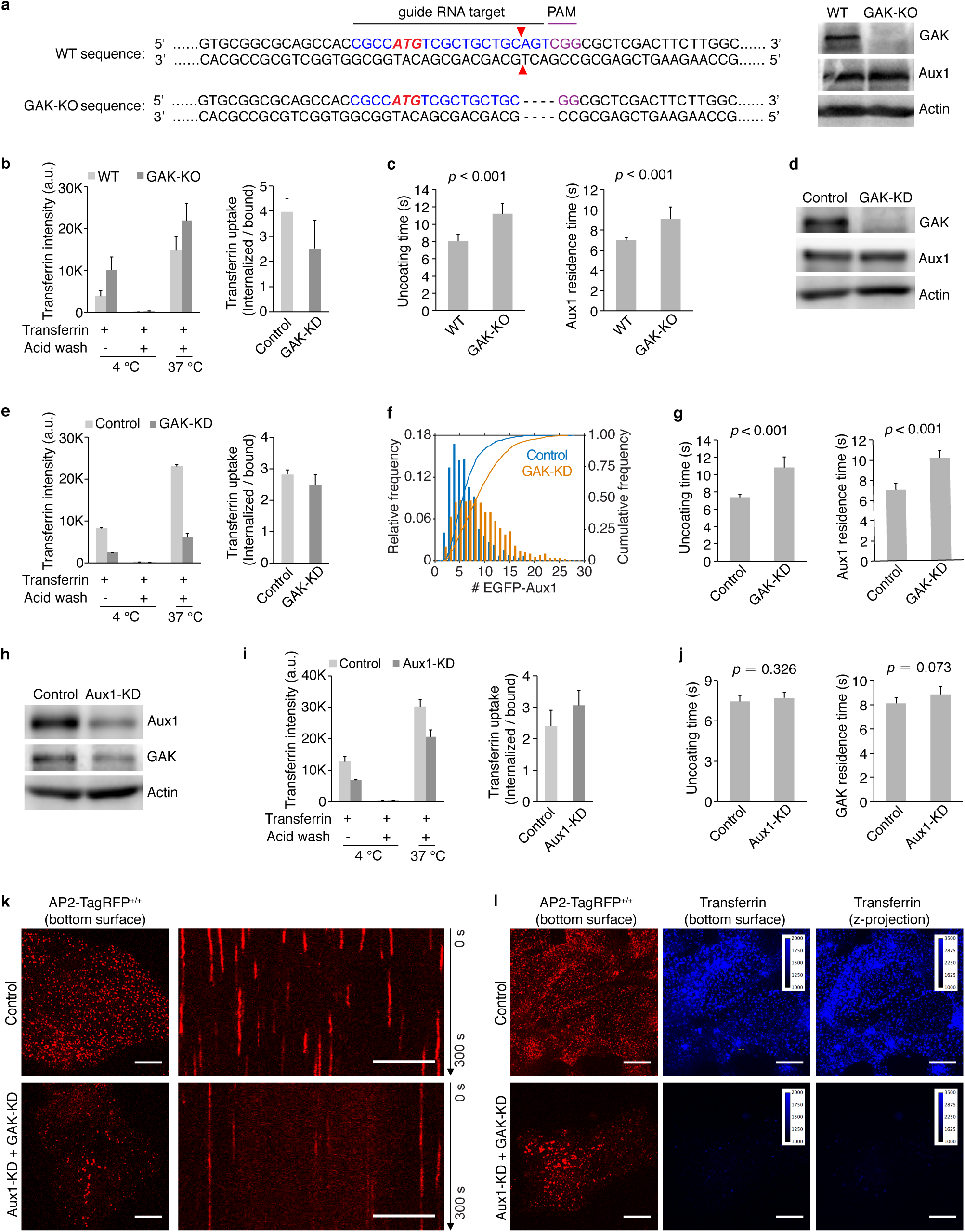
Effects on clathrin-mediated endocytosis of knockout or knockdown of GAK and knockdown of Aux1. (**a**) CRISPR/Cas9 gene-editing strategy used to knock out GAK in cells gene-edited for EGFP-Aux1^+/+^ and CTLA-TagRFP^+/+^. The double strand break (red triangles) induced by Cas9 resulted in elimination of four nucleotides (dotted lines). Loss of GAK expression was confirmed by western blot with antibodies against GAK, Aux1/GAK and actin (right panel). (**b**) Effect of GAK knockout on receptor-mediated uptake of transferrin (n = 3 experiments, mean ± S.D.). (**c**) Uncoating time and Aux1 residence time, in cells lacking GAK. Data from bottom surfaces of double gene-edited EGFP-Aux1^+/+^ and CLTA-TagRFP^+/+^ cells with GAK (n= 5 cells) or lacking GAK by knockout (n = 7 cells) imaged at 1 s intervals for 200 s by TIRF microscopy (mean ± S.D., P values by two-tailed *t*-test). (**d**) Western blot analysis of EGFP-Aux1^+/+^ and CLTA-TagRFP^+/+^ cells treated with lentivirus containing control shRNA (Control) or shRNA specific for GAK (GAK-KD), showing specific reduction of GAK expression 5 days after transduction. (**e**) Effect of GAK knockdown on receptor-mediated uptake of transferrin (n = 2 experiments, mean ± S.D.). See legend for panel (**b**). (**f**) Influence of GAK depletion on Aux1 recruitment. Data from bottom surfaces of double gene-edited EGFP-Aux1^+/+^ and CLTA-TagRFP^+/+^ cells with GAK (1058 traces, 8 cells) or depleted of GAK by knockdown (1380 traces, 9 cells) imaged at 1 s intervals for 200 s by TIRF microscopy. The number of recruited EGFP-Aux1 molecules is significantly increased (Cohen’s *d* = 0.68). (**g**) Influence of GAK depletion on uncoating time (left) and Aux1 residence time (right). Data from bottom surfaces of double gene-edited EGFP-Aux1^+/+^ and CLTA-TagRFP^+/+^ cells with GAK (n = 5 cells) or depleted of GAK by knockdown (n = 5 cells) imaged at 1 s intervals for 200 s by TIRF microscopy (mean ± S.D., P values by two-tailed *t*-test). (**h**) Western blot analysis of parental SUM159 cells incubated with lentivirus containing control shRNA or shRNA specific for Aux1 (Aux1-KD) showing specific reduction of Aux1 expression 5 days after transduction. (**i**) Effect of Aux1 knockdown in gene-edited EGFP-GAK^+/+^ and CTLA-TagRFP^+/+^ cells on receptor-mediated uptake of transferrin (n = 2 experiments, mean ± S.D.). (**j**) Influence of Aux1 depletion on uncoating time (left) and GAK residence time (right). Data from bottom surfaces of double gene-edited EGFP-GAK^+/+^ and CLTA-TagRFP^+/+^ cells with Aux1 (n = 5 cells) or depleted of Aux1 by knockdown (n = 5 cells) imaged at 1 s intervals for 200 s by TIRF microscopy (mean ± S.D., P values by two-tailed *t*-test). (**k**) Bottom surfaces of AP2-TagRFP^+/+^ cells treated with lentivirus containing control shRNA or a mixture of shRNA targeting Aux1 and GAK (Aux1-KD + GAK-KD) imaged at 2 s intervals for 300 s by spinning-disk confocal microscopy. The representative images are from a single time point; the corresponding kymograph shows the entire time series. Scale bars, 10 μm. (**l**) AP2-TagRFP^+/+^ cells with or without double Aux1+GAK knockdown incubated with 10 μg/ml Alexa Fluor 647-conjugated transferrin for 10 min at 37°C and then imaged in 3D using spinning-disk confocal microscopy (30 imaging planes spaced 0.35 μm). Images from the bottom surface of control cells show diffraction-limited AP2-TagRFP spots associated with endocytic coated pits and coated vesicles; in cells depleted of Aux1 and GAK, the punctate distribution is replaced by characteristic larger patches. The images also show the extent of surface binding (bottom surface) and internalization (maximum z-projection of the 30 stacks) of transferrin in the control cells and its absence in the cells impaired in endocytosis due to the Aux1 and GAK depletion. Scale bars, 10 μm.

**Supplementary Figure 5.**
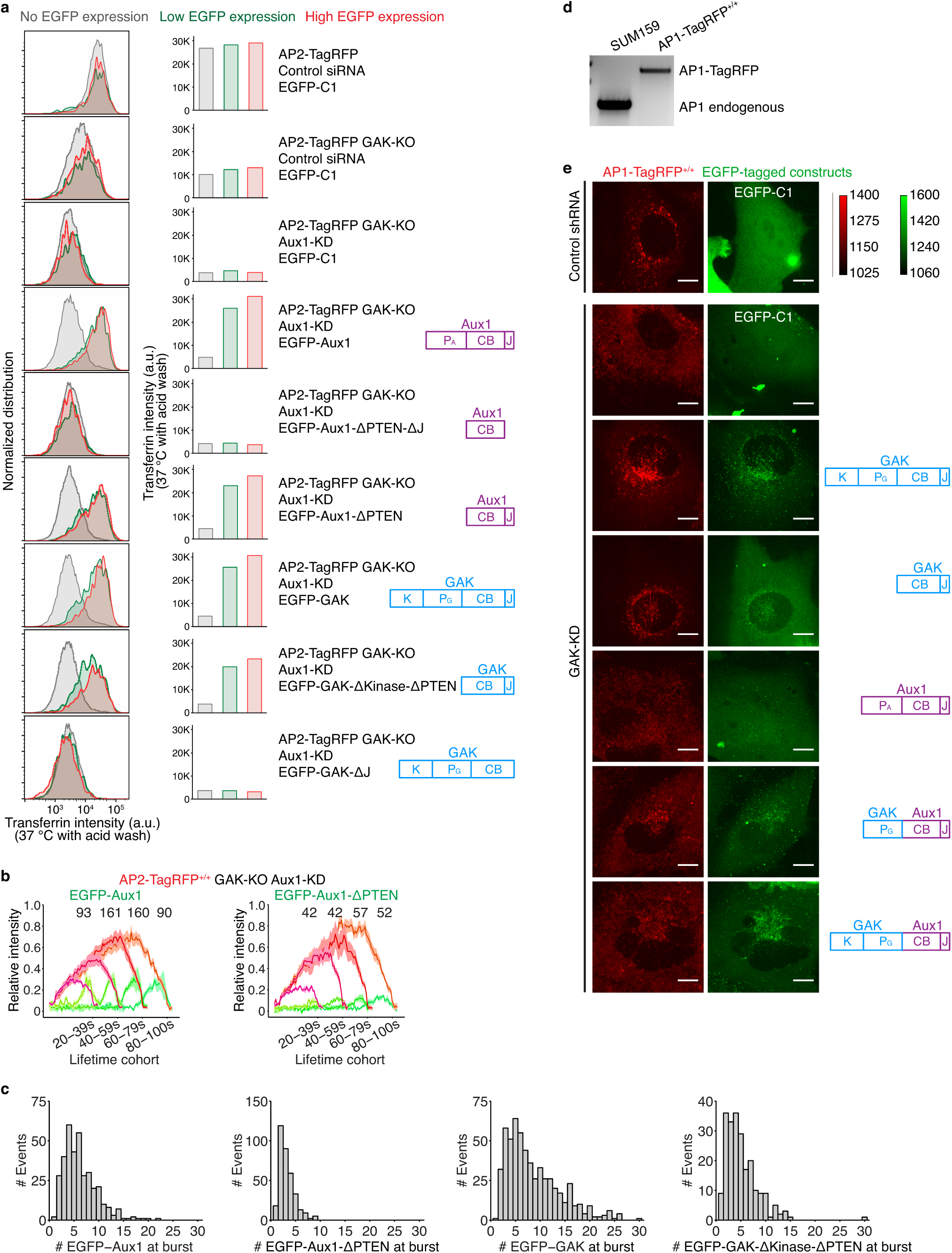
Roles of the PTEN-like domain and clathrin-binding domain of auxilins in the endocytic and secretory pathways. (**a**) AP2-TagRFP^+/+^ cells with or without GAK (AP2-TagRFP GAK-KO) treated with control siRNA or siRNA targeting Aux1 for 3 days (2 sequential transfections), then subjected to transient expression of the indicated EGFP-tagged constructs for additional 1 days followed by measurements of Alexa Fluor 647-conjugated transferrin uptake by flow cytometry. The plots (left panels) and equivalent histograms (right panels) show comparisons of the internalized transferrin (37°C with acid wash) in the absence or presence of low and high levels of ectopic expression of the indicated constructs. (**b**) The GAK-KO AP2-TagRFP^+/+^ cells were treated with siRNA targeting endogenous Aux1 and then transfected for transient expression of EGFP-tagged full length Aux1 (left panel) or Aux1 lacking the PTEN-like domain (right panel). The cells were imaged at 1 s intervals for 300 s by TIRF microscopy. The averaged fluorescence intensity traces (mean ± S.E.) for AP2-TagRFP (red) and EGFP-tagged constructs (green) were identified in 9 and 7 cells, respectively, and then grouped in cohorts according to lifetimes. The numbers of analyzed traces are shown above each cohort. (**c**) The GAK-KO AP2-TagRFP^+/+^ cells were treated with siRNA targeting endogenous Aux1 and then transfected for transient expression of EGFP-tagged constructs as indicated. The cells with EGFP expression at a similar level as the endogenous auxilins were imaged at 1 s intervals for 300 s by TIRF microscopy. Distribution of the maximum number of EGFP-tagged molecules recruited during the uncoating burst (From the left to the right panel: 363 traces from 5 cells, 348 traces from 6 cells, 587 traces from 5 cells, 221 traces from 5 cells, respectively). (**d**) Genomic PCR analysis showing biallelic integration of TagRFP into the *AP1S1* genomic locus to generate the clonal gene-edited cell line AP1-TagRFP^+/+^. (**e**) AP1-TagRFP^+/+^ cells treated with lentivirus containing control shRNA or shRNA targeting GAK for 4 days, then subjected to transient expression of the indicated EGFP-tagged constructs for additional 1 day, and volumetrically imaged by spinning-disk confocal microscopy (34 sequential optical sections spaced 0.3 μm). Maximum intensity z projections acquired using the same acquisition parameters as with gene-edited EGFP-GAK^+/+^ cells, making it possible to identify cells ectopically expressing at the same level as endogenous GAK. Scale bars, 10 μm.

**Supplementary Figure 6.**
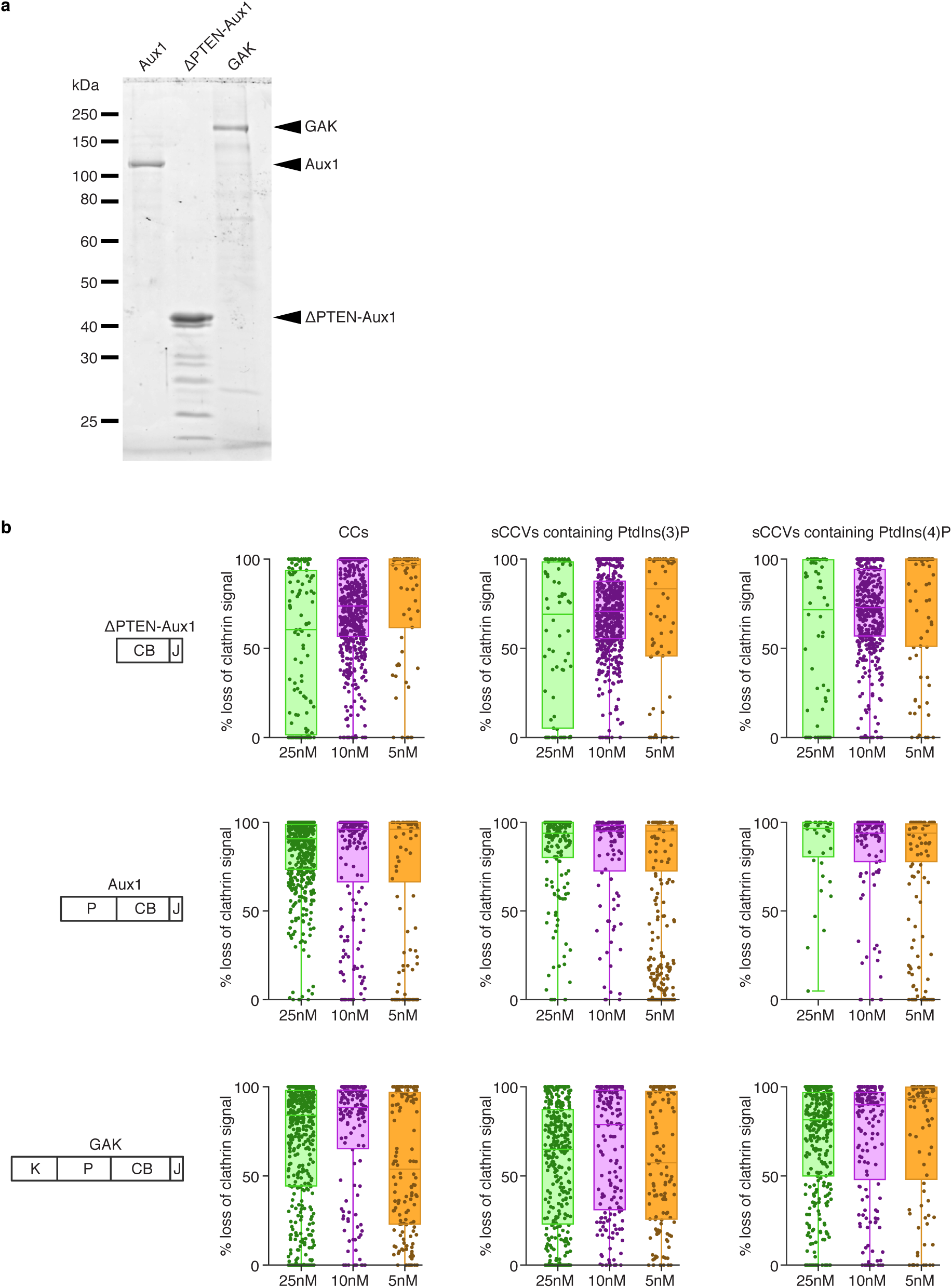
*In vitro* disassembly of clathrin/AP2 coats and synthetic clathrin/AP2 coated vesicles containing PtdIns(3)P or PtdIns(4)P. (**a**) SDS-PAGE (and Coomassie Blue staining) of the recombinant full length Aux1, ΔPTEN-Aux1 and full length GAK. Molecular weight markers are shown. For GAK and ΔPTEN-Aux1, impurities (of high electrophoretic mobility relative to the target species) reduced the full-length target protein proportion to 60% and 50%, respectively (estimated by band densitometry). (**b**) Single-object uncoating efficiency determined from the loss of the clathrin LCa-Alexa Fluor 488 fluorescence signal as a function of ΔPTEN-Aux1, full length Aux1 or full length GAK concentration (5-25 nM range) added together with 1 µM Hsc70 and 5 mM ATP. Each sample included a mixture of clathrin/AP2 coats (CC) together with synthetic clathrin/AP2 coated vesicles containing PtdIns(4,5)P_2_ together with either PtdIns(3)P or PtdIns(4)P (distinguished by labeling with DiI or DiD lipid dyes). Data was acquired at 1 s intervals for 150 s using 3-color TIRF microscopy; each dot in the box plots represents the final uncoating efficiency for a single object. Box plots include the median and data are from three independent experiments.

## Supplementary Video Legends

**Supplementary Video 1. Dynamics of EGFP-Aux1 recruitment to clathrin-coated vesicles.** Bottom surface of a SUM159 cell gene-edited for EGFP-Aux1^+/+^ and CLTA-TagRFP^+/+^ was imaged by TIRF microscopy every 1 s for 200 s. To facilitate visualization, the EGFP channel was shifted laterally by 5 pixels in the right panel.

**Supplementary Video 2. Dynamics of EGFP-GAK recruitment to clathrin-coated vesicles.** Bottom surface of a SUM159 cell gene-edited for EGFP-GAK^+/+^ and CLTA-TagRFP^+/+^ was imaged by TIRF microscopy every 1 s for 200 s. To facilitate visualization, the EGFP channel was shifted laterally by 5 pixels in the right panel.

**Supplementary Video 3. Sequential recruitment of EGFP-Aux1 and TagRFP-GAK to clathrin-coated vesicles.** Bottom surface of a SUM159 cell gene-edited for EGFP-Aux1^+/+^ and TagRFP-GAK^+/+^ was imaged by TIRF microscopy every 1 s for 120 s. To facilitate visualization, the TagRFP channel was shifted laterally by 5 pixels in the right panel.

**Supplementary Video 4. Single-object *in vitro* disassembly of clathrin/AP2 coats and synthetic clathrin/AP2 coated vesicles.** The still image imaged with TIRF at the beginning of the time series corresponds to clathrin/AP2 coats and synthetic clathrin/AP2 coated vesicles containing PtdIns(4,5)P_2_ together with either PtdIns(3)P or PtdIns(4)P distinguished by labeling with DiI (red) or DiD (blue) lipid dyes. Prior to the time series, the channels corresponding to clathrin LCa-Alexa Fluor 488 (green) were shifted 5 pixels with respect to the lipids. The time series follows the uncoating reaction and corresponds to the clathrin fluorescence signal as a function of 25 nM ΔPTEN-Aux1 added together with 1 µM Hsc70 and 5 mM ATP. Data was acquired at 1 s intervals for 150 s using 3-color TIRF microscopy.

